# Dendrite architecture determines mitochondrial distribution patterns *in vivo*

**DOI:** 10.1101/2022.07.01.497972

**Authors:** Eavan J. Donovan, Anamika Agrawal, Nicole Liberman, Jordan I. Kalai, Nicholas J. Chua, Elena F. Koslover, Erin L. Barnhart

## Abstract

Mitochondria are critical for neuronal function and must be reliably distributed through complex neuronal architectures. By quantifying *in vivo* mitochondrial transport and localization patterns in the dendrites of *Drosophila* visual system neurons, we show that mitochondria make up a dynamic system at steady-state, with significant transport of individual mitochondria within a stable global pattern. Mitochondrial motility patterns are unaffected by visual input, suggesting that neuronal activity does not directly regulate mitochondrial localization *in vivo*. Instead, we present a mathematical model in which four simple scaling rules enable the robust self-organization of the mitochondrial population. Experimental measurements of dendrite morphology validate key model predictions: to maintain equitable distribution of mitochondria across asymmetrically branched subtrees, dendritic branch points obey a parent-daughter power law that preserves cross-sectional area, and thicker trunks support proportionally bushier subtrees. Altogether, we propose that “housekeeping” requirements, including the need to maintain steady-state mitochondrial distributions, impose constraints on neuronal architecture.

## INTRODUCTION

Neurons are energetically demanding cells with complex morphologies (Cajal, 1888; Attwell and Laughlin, 2001; Harris et al., 2012). Mitochondria produce most of the ATP in neurons (Hall et al., 2012) and must be distributed throughout all neuronal compartments to meet energetic demands. Mitochondria are also highly dynamic: they grow and degrade, divide and fuse, and move through the cell (Morris and Hollenbeck, 1995; Amiri and Hollenbeck, 2008; Plucinska et al., 2012; Cagalinec et al., 2013; Ashrafi et al., 2014; Misgeld and Schwarz, 2017; Lewis et al., 2018). Thus, to maintain stable mitochondrial distribution patterns over time, neurons must coordinate mitochondrial dynamics over large, elaborately branched axonal and dendritic arbors.

Mitochondrial motility, in particular, has been extensively studied in neurons (Morris and Hollenbeck, 1995; Pilling et al., 2006; Wang et al., 2011; Plucinska et al., 2012; Lewis et al., 2016; Vagnoni and Bullock, 2018; Mandal and Drerup, 2019). For an individual mitochondrion, the mechanics governing motility are relatively clear (Barnhart, 2016). In brief, the motor proteins kinesin and dynein transport mitochondria along microtubules (Pilling et al., 2006). Adaptor proteins link mitochondria to motor proteins, and anchoring proteins oppose mitochondrial movement (Kang et al., 2008; Pathak et al., 2010; Schwarz, 2013). Whether a particular mitochondrion moves, and in which direction, depends on the number and orientation of microtubule tracks, as well as the relative amount of force generated by populations of motor proteins versus anchoring interactions (Barnhart, 2016). It is unclear, however, how neurons regulate these molecular-scale interactions across space and time in order to maintain large-scale mitochondrial distribution patterns.

In principle, neurons could regulate mitochondrial localization patterns by controlling the relative flux of mitochondria into subcellular compartments. In this scenario, mitochondria would be enriched in regions where more microtubule tracks allow for higher rates of mitochondrial delivery. EM studies have shown that microtubule numbers are proportional to the thickness of neuronal processes in various cell types (Kubota et al., 2011; Katrukha et al., 2021), so the shape of a neuron is likely to play a role in mitochondrial transport patterns. The relationship between transport rates and morphological scaling across branch points in tree-like structures — i.e. the relative thickness of parent and daughter branches, as well as the relative sizes of sister subtrees that sprout from each branch point — has been studied in various contexts, from botanical trees (Leonardo et al., 1970; Eloy, 2011; Lehnebach et al., 2018) to the vascular system (Murray, 1926). In neurons, theoretical work has defined various scaling rules that optimize wiring economy (Cherniak et al., 1999; Wen and Chklovskii, 2008; Cuntz et al., 2010), electrical signaling (Rall, 1959; van Ooyen et al., 2002; Chklovskii and Stepanyants, 2003; Wen and Chklovskii, 2008), and, to a lesser extent, transport (Hillman, 1979; Liao et al., 2021). Proposed rules include Da Vinci’s Rule (Leonardo et al., 1970; Hillman, 1979), Rall’s Law (Rall, 1959), and Murray’s Law (Murray, 1926; Cherniak et al., 1999; Chklovskii and Stepanyants, 2003), which conserve, narrow, or expand the total cross-sectional area of the branches at each junction, respectively. However, there is no comprehensive theory or experimental evidence that defines the relationship between neuronal architecture, mitochondrial transport, and large-scale mitochondrial distribution patterns.

To interrogate the relationship between neuronal architecture and dynamic mitochondrial localization patterns *in vivo*, we used neurons in the *Drosophila* visual system called horizontal system (HS) neurons as a model system. There are three HS neurons per optic lobe of the fly (six in total), all of which are functionally equivalent (Schnell et al., 2010). HS neurons have highly branched dendritic arbors, which detect global optic flow patterns by integrating local motion input signals (Barnhart et al., 2018).

To determine how dendritic branching patterns contribute to steady-state mitochondrial localization patterns in HS cells, we combined experimental measurements of *in vivo* mitochondrial transport, mitochondrial localization patterns, and dendrite architecture with mathematical modeling. We found that although mitochondria in HS dendrites are highly motile, large-scale mitochondrial distribution patterns are conserved across HS cells: mitochondria are consistently enriched in distal dendrites, relative to primary dendrites, and equitably distributed across asymmetric sister subtrees. Our experimental measurements are consistent with a mathematical model in which simple dendritic scaling rules enable robust self-organization of steady-state mitochondrial localization patterns. Consistent with previously published work (Faits et al., 2016; Smit-Rigter et al., 2016; Silva et al., 2021), we also found that physiological neuronal activity does not affect mitochondrial motility. Altogether, this work demonstrates that in HS neurons, dendritic architecture, not neuronal activity, determines mitochondrial localization patterns *in vivo*.

## RESULTS

### Mitochondrial localization patterns in HS neurons

To measure mitochondrial distribution patterns in HS neurons, we took advantage of publicly available serial section transmission electron microscopy (ssTEM) images of an entire fly brain (“Female Adult Fly Brain,” or FAFB (Zheng et al., 2018)). We used existing HS skeletons, traced throughout the three-dimensional FAFB image volume (Michael Reiser, unpublished data), to identify mitochondria within HS neurons (Figure 1A-B, S1A-B). Then, we measured mitochondrial morphology as a function of subcellular compartment by manually reconstructing whole mitochondria within small portions of the axon or dendrite (Figure S1A-B). We found that the median volume of an individual mitochondrion was ~0.5 μm^3^ in both HS dendrites and axons (Figure S1C). We also found that dendrites, but not axons, often contained large, branched mitochondria that spanned multiple dendritic branches (Figure S1C), consistent with previously published measurements of mitochondrial morphology in vertebrate neurons (Popov et al., 2005; Turner et al., 2022).

**Figure 1:**
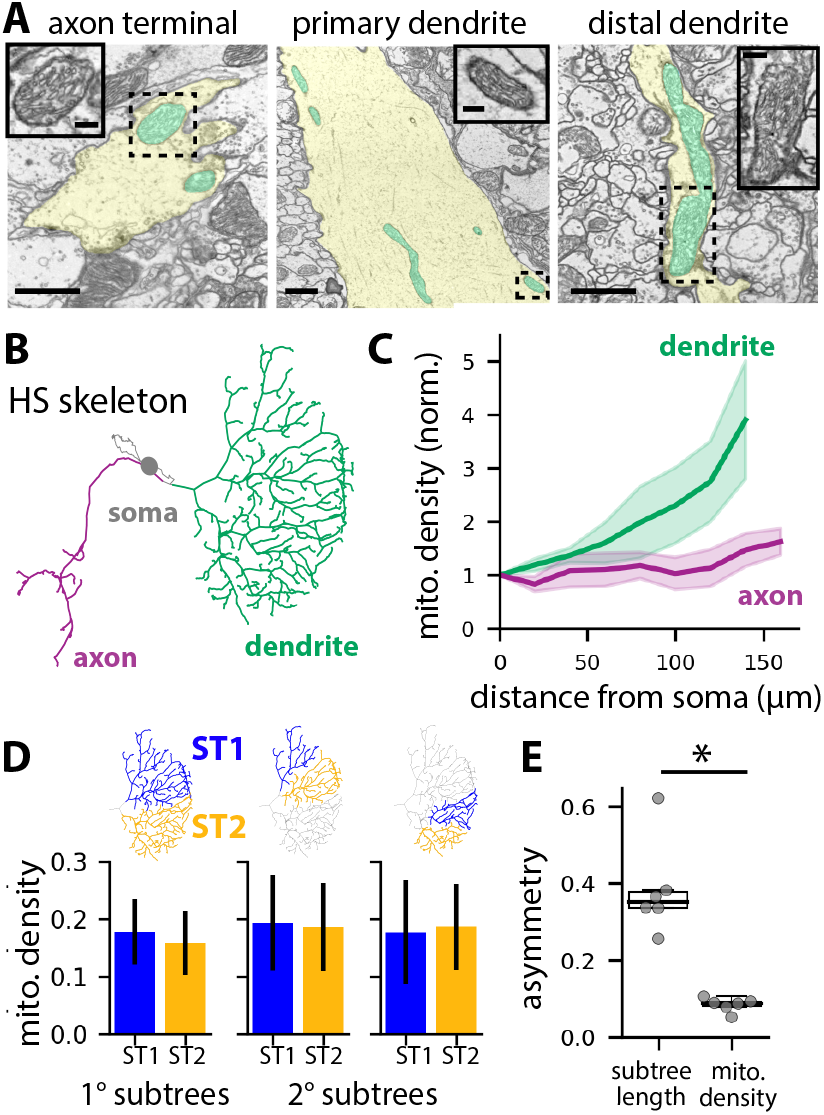
Mitochondrial localization patterns in HS dendrites. A: TEM images of mitochondria (cyan) in different compartments of an HS neuron (yellow); scale bars are 1 μm. Dashed boxes indicate the regions enlarged in the inset images (inset scale bars are 200 nm). B: Skeleton of an HS neuron traced through ssTEM images. C: Average mitochondrial densities plotted versus distance from the soma in the dendrite (green) and axon (magenta). N = 6 neurons; shaded regions indicate the standard error of the mean. For each cell, mitochondrial densities are normalized to the density in the primary dendrite near the soma. D: Mitochondrial densities in sister subtree pairs. Error bars are bootstrap 95% confidence intervals. E: Sister subtree asymmetries in subtree length and mitochondrial density (asymmetry = ((ST1-ST2)^2^/(ST1+ST2)^2^)^1/2^, averaged over all sister subtree pairs per cell); p < 0.01 (T-test).

Next, rather than measuring the size of individual mitochondria within small portions of the cell, we sought to quantify mitochondrial distribution patterns across the entire cell. Manual segmentation of mitochondria in three dimensions, however, is prohibitively labor- and time-intensive. To rapidly measure mitochondrial densities (i.e. the fraction of the neuron occupied by mitochondria) for whole cells, we resampled each HS skeleton such that skeleton nodes were placed at regular 5 μm intervals along the skeleton. Then we extracted two-dimensional image slices centered around each node and manually reconstructed the HS neuronal segment and all mitochondria within it in each image (Figure 1A). We calculated total mitochondrial densities for each HS neuron by dividing the total mitochondrial area by the total neuronal area and found that, on average, the total mitochondrial density in HS neurons is ~20% (Figure S1D; mitochondrial density = 0.19 +/- 0.04 STE). Although we measured significant variation in the total mitochondrial density across the six HS neurons in the FAFB dataset (Figure S1D), we found that mitochondrial localization patterns were conserved across all six cells (Figure 1C,E and S1E-F).

First, we measured mitochondrial densities as a function of subcellular compartment and found consistently higher densities in dendrites than in axons (Figure S1E). Next, in dendrites but not axons, mitochondrial densities increased with distance from the soma, such that mitochondrial densities were approximately four times higher in the distal-most dendrites compared to the primary dendrite (Figure 1C, S1F-G). Finally, we measured mitochondrial densities across dendritic subtrees. At each branch point within a dendritic arbor, a parent branch splits into two daughter branches, and the entire arbor can be decomposed into successive pairs of sister subtrees (Figure 1D). Although in some cases the dendritic arbor splits in a symmetric fashion, we found that sister subtrees are often asymmetric (Figure 1E). Despite this asymmetry in sister subtree size, we found that mitochondria are equitably distributed across branch points such that sister subtrees have equivalent mitochondrial densities (Figure 1D-E). Altogether, these results show that mitochondria in HS dendrites follow a specific distribution pattern, with equitable distribution across sister subtrees and mitochondrial enrichment in distal dendrites.

### *In vivo* mitochondrial motility in HS dendrites

Mitochondria in neurons are motile, both in cell culture (Morris and Hollenbeck, 1995; Overly et al., 1996; Wang and Schwarz, 2009) and *in vivo* (Plucinska et al., 2012; Lewis et al., 2016; Mandal and Drerup, 2019; Silva et al., 2021). We therefore hypothesized that the specific mitochondrial localization pattern we measured in HS dendrites reflects a dynamic steady-state in which individual mitochondria are continually reorganizing within a stable, global pattern. To test this hypothesis, we used the GAL4/UAS binary system to drive expression of GFP-tagged mitochondria (mitoGFP) and a cytosolic volume marker (tdTomato) in HS neurons (Brand and Perrimon, 1993). Then, we used *in vivo* confocal microscopy to image HS dendrites in living, head-fixed *Drosophila* (Figure 2A). Mitochondria are densely packed within HS dendrites, so we bleached stationary mitochondria in a small portion of the dendrite to resolve motile mitochondria moving through the bleached region. We measured mitochondrial motility in primary and distal dendritic branches (Figure 2B-D). In the primary dendrite, mitochondria moved in both anterograde (into the arbor) and retrograde (out of the arbor) directions (Figure 2D). Motile mitochondria were highly persistent, with less than 20% of mitochondria arresting or pausing as they moved through the primary dendrite (Figure S2A). Individual mitochondria exhibited a broad range of speeds and lengths, but there were no significant differences between mitochondria moving in the anterograde versus retrograde directions in the primary dendrite (Figure 2E-F). In the distal dendrites, mitochondria moving in the anterograde direction were significantly slower than in the primary dendrite (Figure 2E). Linear flux rates were also significantly lower in the distal dendrites (Figure S2B). Finally, the average flux of anterograde mitochondria moving through the primary dendrite into the arbor was equivalent to the average flux of retrograde mitochondria moving back out of the arbor (Figure 2G). This balanced anterograde and retrograde transport through the primary dendrite suggests that the total amount of mitochondria in the HS arbor remains constant over time.

**Figure 2:**
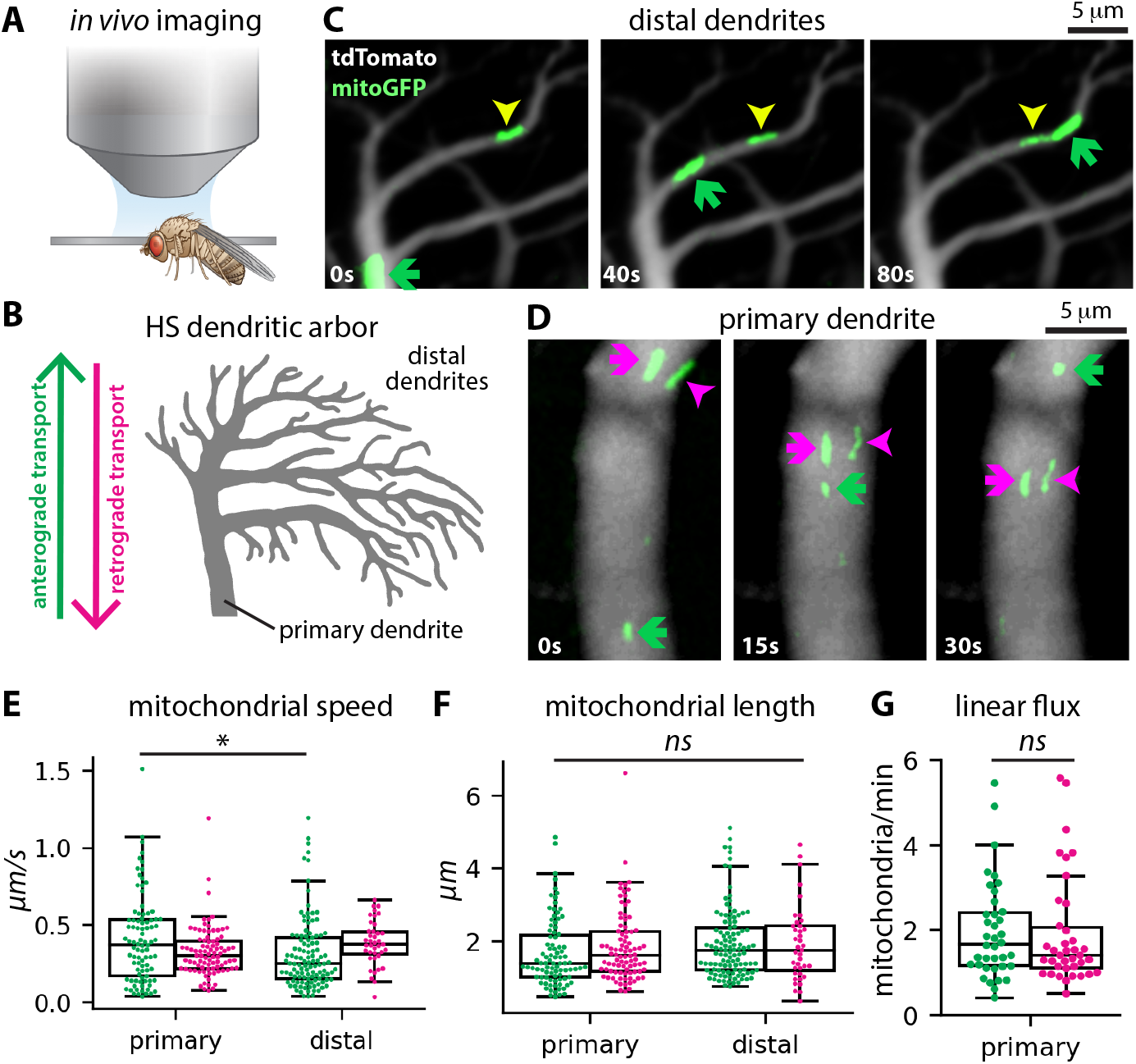
Mitochondrial transport in HS dendrites. A: Experimental setup with confocal microscope positioned above head-fixed fly. B: Schematic depicting an HS dendritic arbor, including primary and distal dendrites and the direction of anterograde versus retrograde transport. C-D: Image time series showing mitochondrial transport in distal (C) and primary (D) dendrites. Arrows indicate mitochondria moving in the anterograde (green) and retrograde (magenta) directions, along with an arrested mitochondrion (yellow). E-F: Average speed (E) and length (F) of mitochondria moving through primary or distal mitochondria in the anterograde (green) or retrograde (magenta) directions. Each dot represents an individual mitochondrion. The asterisk indicates a significant difference (one way ANOVA, post-hoc Tukey’s test). G: Mitochondrial linear flux rates (number of mitochondria per minute) in primary dendrites in the anterograde (green) and retrograde (magenta) directions. Dots represent individual primary dendrites. There is no significant difference in anterograde versus retrograde linear flux (T-test).

Notably, our measurements of mitochondrial density and motility allow us to estimate the fraction of dendritic mitochondrial volume that is motile versus stationary at any given point in time. In the primary dendrite, approximately two mitochondria pass a cross-section per minute, in each direction. Based on our measurements of mitochondrial length, we estimate that the volume of each motile mitochondrion is ~0.5 μm^3^ (see Methods), consistent with the median mitochondrial volume in our EM reconstructions (Figure S1C). Thus, ~1 μm^3^ of mitochondrial mass exchanges through the primary dendrite every minute. Based on this rate of mitochondrial volume exchange (J ~1 μm^3^/min), the typical speed of motile mitochondria (v ~0.4 μm/s), the mitochondrial volume density (c ~10%), and the cross-sectional area (A_d_ ~ 30 μm^2^) in the primary dendrite, we estimate that the fraction of motile mitochondria mass in this proximal region is f_m_ = 2 J/(v c A_d_) ~ 3%. Mitochondrial density is higher in the distal dendrites (c ~30%), and we therefore estimate an even lower motile fraction for the distal portions of the dendritic arbor (f_m_ ~1%). Thus, the majority of the mitochondrial population is stationary at any given moment. However, we also estimate that the total volume of mitochondria in the dendritic arbor exchanges through the primary dendrite many times over the lifetime of the neuron. Based on the mitochondrial density in the whole dendrite (c ~20%), the total volume of the dendrite (V_d_ ~2000 μm^3^ (Cuntz et al., 2013) and the rate of mitochondrial volume exchange (J ~ 1 μm^3^/min), we estimate the fraction of mitochondrial volume exchanged per unit time in the dendrite to be: J_norm_ = J/(c V_d_) ~15% hr^−1^. At this rate, the entire mitochondrial volume in an HS arbor reorganizes in less than ten hours, or more than one hundred times over the course of a fly’s lifetime. Altogether, these results are consistent with the idea that the mitochondria in HS dendrites make up a dynamic system at steady-state.

### Visual stimulus-evoked calcium signals do not affect mitochondrial motility in HS dendrites

How do HS dendrites maintain steady-state mitochondrial distribution patterns, given rapid reorganization of mitochondrial mass? HS neurons selectively respond to specific patterns of global optic flow (Schnell et al., 2010; Fujiwara et al., 2017; Barnhart et al., 2018), which drive large calcium increases in the distal dendrites of HS neurons (Barnhart et al., 2018). Calcium arrests mitochondrial motility in cultured neurons (MacAskill et al., 2009; Wang and Schwarz, 2009), so we reasoned that if stimulus-evoked calcium signals are stronger in the distal dendrites than in the primary dendrite, then calcium-dependent arrest of mitochondrial transport could increase the density of mitochondria in the distal dendrites, relative to the primary dendrite. To test this idea, we projected the preferred visual stimulus for HS neurons — a global motion stimulus moving from front to back across one eye — on a screen positioned in front of the fly while simultaneously imaging a genetically-encoded calcium reporter (RGECO1a) in HS dendrites (Figure 3A, S3A). Consistent with our previously published work, front-to-back global motion stimuli drove robust calcium responses in HS distal dendrites (Figure S3A-C). In contrast, we measured significantly smaller calcium responses to the same stimuli in primary dendrites (Figure S3A-C). Thus, stimulus-evoked calcium signals correlate with mitochondrial density, with larger calcium response amplitudes and mitochondrial densities in the distal dendrites compared to the primary dendrites.

**Figure 3:**
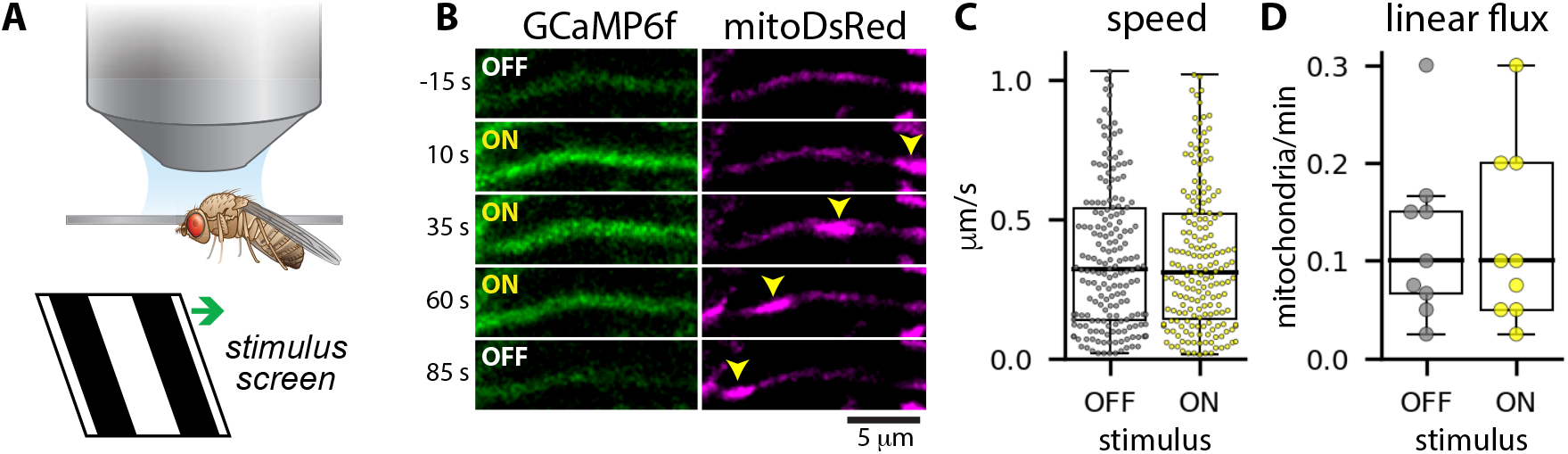
Stimulus-evoked calcium signals do not affect mitochondrial motility in HS dendrites. A: Experimental setup. Neurons are imaged by confocal microscopy while a visual stimulus is presented on a screen in front of the fly. The visual stimulus is squave wave gratings moving in the preferred direction (front-to-back across one eye) for HS neurons. B: Calcium responses to the motion stimulus (GCaMP6f, left) and mitochondria (mitoDsRed, right) in an HS dendritic branch. Yellow arrows indicate a moving mitochondrion. C-D: Mitochondrial speeds (C) and linear flux rates (D) when the visual stimulus was OFF (gray) versus ON (yellow). Dots indicate instantaneous speeds measured between successive frames (C) or average linear flux rates for individual flies (D).

However, we also found that visual stimulus-evoked calcium signals are not sufficient to arrest mitochondrial motility in HS distal dendrites (Figure 3B-D). When we simultaneously imaged a green calcium reporter (GCaMP6f) and mitochondrially-targeted DsRed (mitoDsRed) while presenting global motion stimuli to the fly, we again measured large, stimulus-evoked calcium signals in the distal dendrites (Figure 3B). We observed mitochondria moving through distal dendritic branches, regardless of whether the stimulus was on or off (Figure 3B). The stimulus did not affect mitochondrial speeds (Figure 3C), nor did we measure a difference in mitochondrial transport rates when the stimulus was on versus off (Figure 3D). Thus, physiologically-relevant calcium signals do not directly affect mitochondrial motility in HS dendrites *in vivo*.

### Simple scaling rules recapitulate experimental measurements of mitochondrial localization patterns

#### A mean field model of mitochondrial distributions in branched dendrites

We next hypothesized that dendritic architecture, rather than neuronal activity, determines steady-state mitochondrial localization patterns in HS cells. To test this idea, we developed a quantitative model of mitochondrial transport and localization patterns in HS dendrites. In developing this model, we assumed that mitochondrial distributions are governed by the local relationship between mitochondrial transport rates and dendrite thickness as well as the morphology of the dendritic arbor. Specifically, we defined four simple scaling relationships governing mitochondrial localization patterns (Figure 4A):

1. *Scaling of mitochondrial transport with dendrite radius*. Our *in vivo* measurements of mitochondrial transport show that mitochondria are significantly more likely to arrest motility in thin distal dendrites than in the thick primary dendrite (Figure S2A). Based on this, we assume that the rate of mitochondrial arrest, k_s_, scales with dendrite radius according to k_s_ ~ 1/r^β^. This relationship incorporates a width-dependence for the population-averaged speed of mitochondrial transport in different dendritic branches.
2. *Splitting of mitochondria at branch points*. Motor proteins transport mitochondria along microtubules in neurons (Pilling et al., 2006), and microtubule numbers scale with branch thickness in other neuronal cell types (Kubota et al., 2011; Katrukha et al., 2021). We therefore assume that the probability of a mitochondrion moving into a given daughter branch is proportional to the cross-sectional area of the branch, according to p_1_/p_2_ = r_1_^2^/r_2_^2^, where p_1_ and p_2_ are the probabilities that a mitochondrion moves from the parent into daughter 1 or 2, and r_1_ and r_2_ are the radii of the two daughters.
3. *Power law scaling of parent and daughter branch widths*. Consistent with several previous studies (Hillman, 1979; Cherniak et al., 1999; van Ooyen et al., 2002; Chklovskii and Stepanyants, 2003), we assume that parent and daughter radii scale according to the power law r_0_^α^ = r_1_^α^ + r_2_^α^, where r_0_ is the radius of the parent branch and r_1_ and r_2_ are the radii of the daughter branches. The exponent α determines the extent to which the total dendritic cross-sectional area increases or decreases in the daughter branches, relative to the parent: area is conserved for α = 2, decreases for α < 2, and increases for α > 2. Optimal values for α have been derived based on theoretical arguments for preservation of graded electrical signals across dendritic branch points (α = 3/2, often called “Rall’s Law” after the neuroscientist Wilfrid Rall (Rall, 1959)), action potential propagation in axons (α = *3*, often called “Murray’s Law” and first derived for the vasculature system) (Murray, 1926; Cherniak et al., 1999; Chklovskii and Stepanyants, 2003; Mandal and Drerup, 2019), or efficient intracellular transport (α = 2, often called “Da Vinci scaling” after Da Vinci’s rule for trees) (Leonardo et al., 1970; Hillman, 1979).
4. *Scaling of sister subtree size with trunk thickness*. Finally, we assume that larger subtrees are supported by proportionally thicker trunks, with trunk radii scaling with either total subtree length, depth, or some other measure of subtree size. This scaling, along with proportional mitochondrial transport at branch points, should ensure that larger subtrees receive a proportionally larger supply of mitochondria.

**Figure 4:**
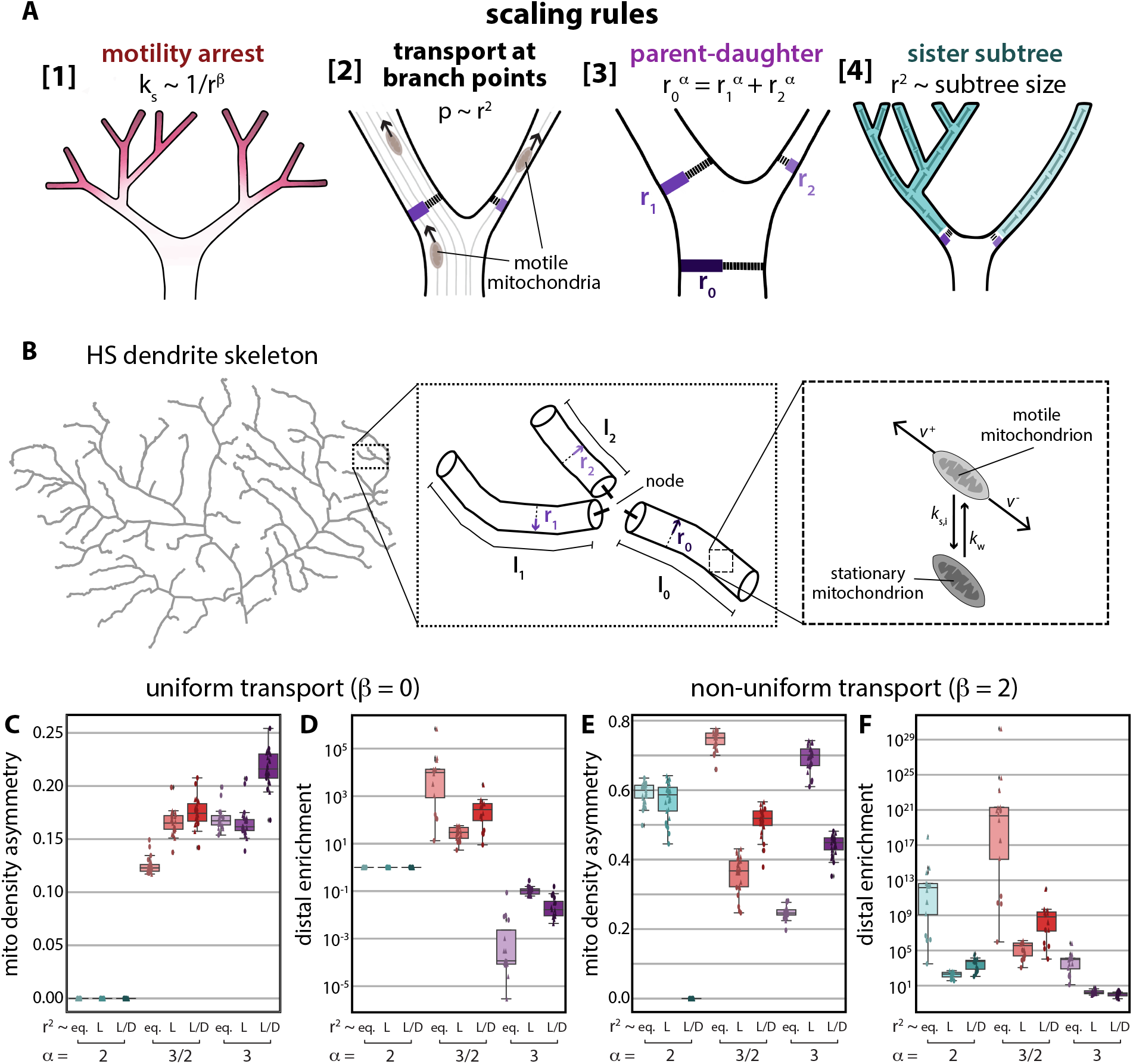
A mean field model for mitochondrial distributions recapitulates experimental measurements when dendrites obey specific scaling rules. A: Scaling rules included in the model. Rule 1: Mitochondrial arrest rates scale with dendrite radius according to k_s_ ~1/r^β^. Rule 2: Mitochondria split at branch points according to the cross-sectional area of each daughter branch. Rule 3: Parent (r_0_) and daughter (r_1_ and r_2_) radii scale according to r_0_^α^ = r_1_^α^ + r_2_^α^. Rule 4: Sister subtrees scale such that thicker trunks support proportionally larger subtrees. Subtree size can be subtree volume, length, or bushiness, where bushiness is length/depth. B: In the model, dendrites are binary trees in which each junction node connects a parent edge to two daughter edges (center panel). Each edge is a cylinder with fixed radius (r) along its entire length (l). The connectivity and length of each edge are extracted from real HS dendrite skeletons (left panel); radii are set according to various forms of parent-daughter and sister-subtree scaling rules (Rules 3 and 4). Within this dendritic structure, mitochondria can move persistently in either the anterograde (v^+^) or retrograde (v^−^) direction, as well as arresting and initiating motility (right panel). For simplicity, we assume a constant motility initiation rate (k_w_) and constant velocities v^+^ and v^−^; arrest rates vary according to k_s_~1/r^β^ (Rule 1). At branch points, motile mitochondria move into daughter branches according to their cross-sectional areas (Rule 2). C-F: Model results. Dendrite topologies (edge length and connectivity for each dendritic branch) were extracted from HS skeletons traced through ssTEM images (circles, N = 6 dendrites, Michael Reiser, unpublished results) or previously published HS dendrite reconstructions (squares, N = 20 dendrites, Cuntz et al 2013). Mitochondrial density asymmetry across sister subtrees (C,E) and distal mitochondrial enrichment (D,F) were calculated for β = 0 (C,D) or β = 2 (E,F), α = 2 (cyan), 3/2 (red), or 3 (purple), and sister subtree scaling with subtree trunks splitting according to r_1_=r_2_ (eq.), r^2^~L, or r^2^~L/D. The mitochondrial density asymmetry is the root-mean squared asymmetry across sister subtrees ST1 and ST2, where asymmetry = (ST1-ST2)/(ST1+ST2). Distal enrichment is mitochondrial density in the distal-most dendritic branches, defined as branches with path distance from the root node >= 75% of the maximum path distance, divided by the mitochondrial density in the primary dendrite. Box plots show the median, interquartile range, and 1.5x the interquartile range.

To determine how these scaling rules contribute to steady-state mitochondrial localization patterns, we developed a mathematical model of mitochondrial transport in dendritic trees. In this model, each dendrite is a binary tree, with each junction node connecting a parent edge to two daughter edges (Figure 4B). We assume that each edge is a cylinder with fixed radius *ri* along its entire length *l_i_*. In our model, the connectivity and length of each edge in a tree are known attributes, either extracted from skeletons of real HS neurons (Figure 4B) or from synthetic trees with varying branching patterns (Figure S4A). Radii of each branch are set according to various rules for parent-daughter and sister subtree scaling (Rules 3 and 4, Figure 4A).

Within this dendritic structure, we assume that discrete mitochondrial units undergo active transport, which includes processive motion in the anterograde and retrograde directions, as well as arrest and restarting (Figure 4B). The ratio of arrest and restarting rates (k_s_/k_w_) determines the local fraction of stationary mitochondria in each dendritic branch. We assume the restarting rate is constant throughout the arbor, but allow the arrest rate to vary in dendrites of different widths (Rule 1, Figure 4A). Finally, we assume that the anterograde mitochondrial linear flux entering a junction node is split in proportion to the cross-sectional area of the daughter branches, so that wider daughter branches receive proportionally more mitochondria (Rule 2, Figure 4A),

We implemented several versions of this model, using different forms of parent-daughter, sister subtree, and transport scaling rules, through analytical solutions of mean-field equations for mitochondrial density (Figures 4, 7, S4, and S7) and stochastic simulations (Figure S5). For all model versions, we calculated mitochondrial densities as a function of distance from the soma and across sister subtrees to determine which scaling rules recapitulate our experimental measurements (i.e. distal enrichment and equitable distribution of mitochondria across sister subtrees).

#### Model results with uniform transport parameters

In the first, simplest version of our model, we assumed that mitochondrial arrest rates and transport speeds are independent of dendrite thickness (*β* = 0 in Rule 1), such that the fraction of stationary versus motile mitochondria is constant throughout the arbor, and that the linear flux of mitochondria splits in proportion to daughter branch thickness at each branch point (Rule 2). We also assumed that parent and daughter radii follow power law scaling (Rule 3), either Rall’s law (α = 3/2), Da Vinci’s rule (α = 2), or Murray’s law (α = 3). When α = 2, the total cross-sectional area is conserved at each junction, giving rise to constant mitochondrial volume densities throughout the entire arbor, regardless of the tree topology and the relative thickness of sister radii (as set by various forms of Rule 4). Thus, Da Vinci parent-daughter scaling results in equitable distribution of mitochondria across sister subtrees, but not distal enrichment in this minimal version of our model (Figure 4C,E, S4B,D).

In contrast, when the narrowing of daughter branches is governed by Rall’s Law (α = 3/2), the reduction in total cross-sectional area at each branch point results in distal enrichment of mitochondria (Figure 4E, S4D). When the narrowing of daughter branches is governed by Murray’s Law (α = 3), the expansion in total cross-sectional area at each branch point results in distal dilution of mitochondria (Figure 4E, S4D). In addition, when α = 3/2 or 3 (or any value other than 2), the relative volume density of mitochondria in sister subtrees depends on the relative thicknesses of sister trunks at each branch point (as set by various forms of Rule 4), and different forms of sister subtree scaling all result in significant mitochondrial density asymmetries unless dendritic branching patterns are perfectly symmetric (Figure 4C, S4B). In particular, subtrees with more branch points tend to accumulate higher or lower mitochondrial densities due to the reduction or expansion of dendritic cross-sectional area below each branch points when α = 3/2 or 3, respectively. Moreover, we show that for an arbor obeying Rall’s Law, it is impossible to establish equitable mitochondrial distributions between asymmetric sister subtrees with any single function that sets sister subtree trunk thicknesses based on subtree morphology (Figure S4F and Supplemental Methods). Altogether, when mitochondrial transport parameters are spatially uniform, our model recapitulates either equitable distribution of mitochondrial across sister subtrees (for Da Vinci-scaled dendrites) or distal enrichment of mitochondria (for Rall-scaled dendrites), but not both.

#### Model results with non-uniform transport parameters

Our experimental measurements indicate that in thin distal dendrites, average mitochondrial speeds are lower and arrest rates are higher than in the thick primary dendrite (Figure 2E and S2A), and we reasoned that scaling of mitochondrial transport with dendrite radius could result in increased mitochondrial densities in thin distal dendrites. We therefore updated our initial model such that mitochondrial transport rates scale with dendrite radii. Specifically, we assumed that mitochondrial arrest rates are inversely proportional to dendrite cross-sectional area, *k_s_* ~ 1/*r^2^* (*β* = 2 in Rule 1). For simplicity, we assumed that mitochondrial motility initiation rates and speeds are still spatially uniform; different assumptions about how mitochondrial transport changes with branch thickness (e.g. scaling of mitochondrial speed with dendrite radius) give the same results when the fraction of motile mitochondria is very small (see Supplemental Methods), as we observed experimentally.

In dendrites that follow Da Vinci scaling of parent and daughter branch widths (α = 2 in Rule 3), this model predicts that motile mitochondria densities will be constant throughout the dendrite, while stationary mitochondrial densities will be higher in distal dendrites (Figure 4F, S4E). If we assume that daughter branch radii are split equally (*r_1_* = *r_2_*) at each junction, then asymmetric topologies of sister subtrees lead to asymmetry in the mitochondrial distribution, with more-branched subtrees acquiring a higher mitochondrial density (Figure 4E, S4D).

Next, we investigated whether an alternate relationship between the trunk widths of sister subtrees could restore equitable distribution of mitochondria across subtree pairs. According to our analytical calculations for Da Vinci trees with *k_s_* ~ 1/*r*^2^ (see Supplemental Methods), the ratio of stationary mitochondrial densities in a pair of sister subtrees is given by *c^(s)^_1_/c^(s)^_2_* = (*L_1_/V_1_)/(L_2_/V_2_*), where *c^(s)^* is the density of stationary mitochondria and *L* and *V* are subtree length and volume. Equitable distribution of mitochondria therefore depends on a specific morphological relationship between sister subtrees: the total subtree volume must be proportional to its length, such that *L_1_*/*V_1_* = *L_2_/V_2_*.

For an arbor that follows the Da Vinci scaling law, the preservation of cross-sectional area across branch points means that the total volume of a tree can be expressed as *V* = *r_0_^2^D*, where *r_0_* is the radius of the trunk of the tree and *D* describes the effective depth of the tree. For a tree where all terminal dendritic branch tips are the same distance *d* from the trunk, the effective tree depth is *D = d*. For trees with distal tips at varying distances from the trunk, we compute the effective depth *D* recursively from the branch lengths in each subtree (see Supplemental Methods). Notably, the dependence of subtree volume on the trunk radius implies that for subtrees with different depths, it is not possible to split sister subtree trunks in a way that would allow their cross-sectional area to be proportional to volume. We thus do not consider r^2^ ~ V splitting behaviors in our calculations.

In a tree with Da Vinci parent-daughter scaling, all subtrees will exhibit a volume proportional to their total branch length if and only if the sister subtree trunk areas scale according to the following relation: *r_1_^2^/r_2_^2^* = (*L_1_*/*D_1_*)/(*L_2_*/*D_2_*). The ratio of total length over depth (*L/D*) can be thought of as the “bushiness” of a tree. In trees with no branch points, *L = D* and thus *L/D* = 1. In trees with many branch points (i.e. bushier trees), *L* >> *D* and *L/D* >> 1.

In dendrites that follow Da Vinci parent-daughter scaling and sister subtree scaling with r2~L/D, mitochondria are equitably distributed across subtrees (Figure 4D, S4C). Trees that follow different sister subtree scaling rules (e.g. with radii splitting proportional to subtree length) do not exhibit such equitable distributions across subtrees (Figure 4D, S4C). Regardless of the choice of parent-daughter or sister subtree scaling rule, increased mitochondrial arrest rates in thinner distal branches leads to distal mitochondrial enrichment (Figure 4F). We note that the predicted distal enrichment of mitochondria in these simple models is unrealistically high. This effect can be mitigated by assuming a smaller value for β, such that mitochondrial arrest rates increase less as dendrite radii decrease, or setting a lower limit on dendrite radii (Liao et al., 2021), as discussed below.

Our analytical mean-field calculations are supplemented by stochastic simulations of point-like mitochondria moving through small synthetic dendritic arbors (Supplemental Figure S5). These simulations show average densities similar to those predicted by the mathematical model, although higher asymmetries are found between sister subtrees due to stochastic variation in the simulations. Both simulations and mean-field calculations demonstrate that this model — with dendritic trees obeying Da Vinci parent-daughter scaling, inverse scaling of mitochondrial arrest and dendrite thickness, and sister subtree trunk areas proportional to tree bushiness — successfully recapitulates the key features of experimentally observed mitochondrial distributions: equitable densities between sister subtrees and increased density in distal branches.

### Mitochondria split according to dendrite thickness at asymmetric branch points

Our model rests on the assumption that mitochondria traversing branch points in the anterograde direction split according to the cross-sectional areas of the daughter branches. To test this assumption, we used our *in vivo* imaging setup to measure mitochondrial transport across primary branch points in HS dendrites (Figure 5A). We found, first, that most mitochondria moving through the branch in the anterograde direction moved persistently from the parent into one of the two daughter branches without reversing direction or arresting motility (Table S1). In the retrograde direction, the vast majority of mitochondria moved from a daughter branch into the parent branch, with infrequent reversals or arrests and almost no mitochondria moving from one daughter to the other (Table S1). At each branch point, the population of mitochondria moving in the anterograde direction splits, with some mitochondria moving from the parent branch into one daughter branch and the rest into the other (Figure 5A). By comparing mitochondrial linear flux rates to daughter branch cross-sectional areas (Figure 5B), we found that asymmetric linear flux rates (where more mitochondria are transported into one daughter than the other) correlate with asymmetric daughter branch cross-sectional areas (where one daughter is thicker than the other). Thus, proportionally more mitochondria are transported into thicker daughter branches, consistent with our assumption that mitochondria split at branch points according to daughter branch cross-sectional area.

**Figure 5:**
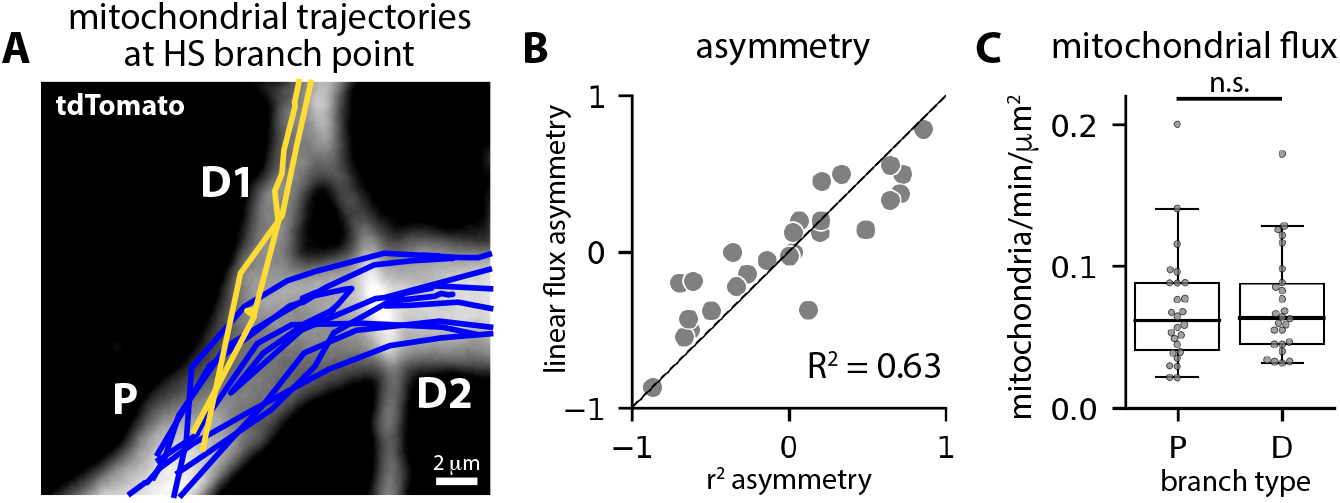
Motile mitochondria split according to dendrite thickness at asymmetric branch points. A: Trajectories of mitochondria moving in the anterograde direction from a parent branch (P) into one of two daughter branches (D1 and D2). Trajectory colors indicate mitochondria that moved into daughter one (D1, yellow) or daughter two (D2, blue). B: Asymmetry in daughter branch radii squared plotted versus asymmetry in mitochondrial linear fiux rates; asymmetry = (D1-D2)/(D1+D2), N = 26 branch points. C: Mitochondrial fiux rates — linear fiux rates (mito-chondria/minute) divided by dendrite cross-sectional area — in parent (P) and daughter (D) branches; n.s. indicates no significant difference (T test).

In addition, we found that mitochondrial flux rates (the linear flux rate divided by branch cross-sectional area) are conserved across branch points, such that flux in the parent branch is roughly equivalent to flux in the daughter branches (Figure 5C, average flux = 0.07 +/- 0.01 (STE) mitochondria/min/um^2^ in both parent and daughter branches). We measured similar mitochondrial flux rates in primary (0.08 +/- 0.01 mitochondria/min/um^2^) and distal (0.06 +/- 0.01 mitochondria/min/um^2^) dendrites as well (Figure S2D). Conservation of mitochondrial flux rates throughout the arbor is consistent with spatially constant motile mitochondrial volume densities which, according to our model, occurs in dendrites that follow Da Vinci parent-daughter scaling.

### HS dendrites follow simple dendritic scaling rules

According to our model, HS dendrites maintain realistic mitochondrial distributions if parent and daughter radii scale according to Da Vinci’s rule (α = 2), and sister subtrees scale with trunk thickness proportional to subtree bushiness (r^2^~L/D). To determine whether HS dendrites obey these morphological scaling rules, we measured HS dendritic architecture. We used stochastic multicolor FlpOut (MCFO) labeling (Nern et al., 2015) to label individual HS dendrites, which we then imaged by confocal microscopy (Figure 6A-B). Next, we segmented and skeletonized each dendrite before measuring parent and daughter branch radii and the length, volume, and bushiness of the subtrees sprouting from each branch point (Figure 6B). We found, first, that HS dendrites are asymmetrically branched, with significant asymmetry in daughter branch thickness and subtree length, volume, and bushiness (Figure 6C). Second, we fit our measurements of parent and daughter radii to the power law r_0_^α^ = r_1_^α^ + r_2_^α^ for a range of values for the exponent α, and we found HS dendrites are approximately Da Vinci-scaled, with α = 2.2 giving the best fit (Figure 6D, R^2^ = 0.87, 95% bootstrap confidence interval = 1.98-2.44).

**Figure 6:**
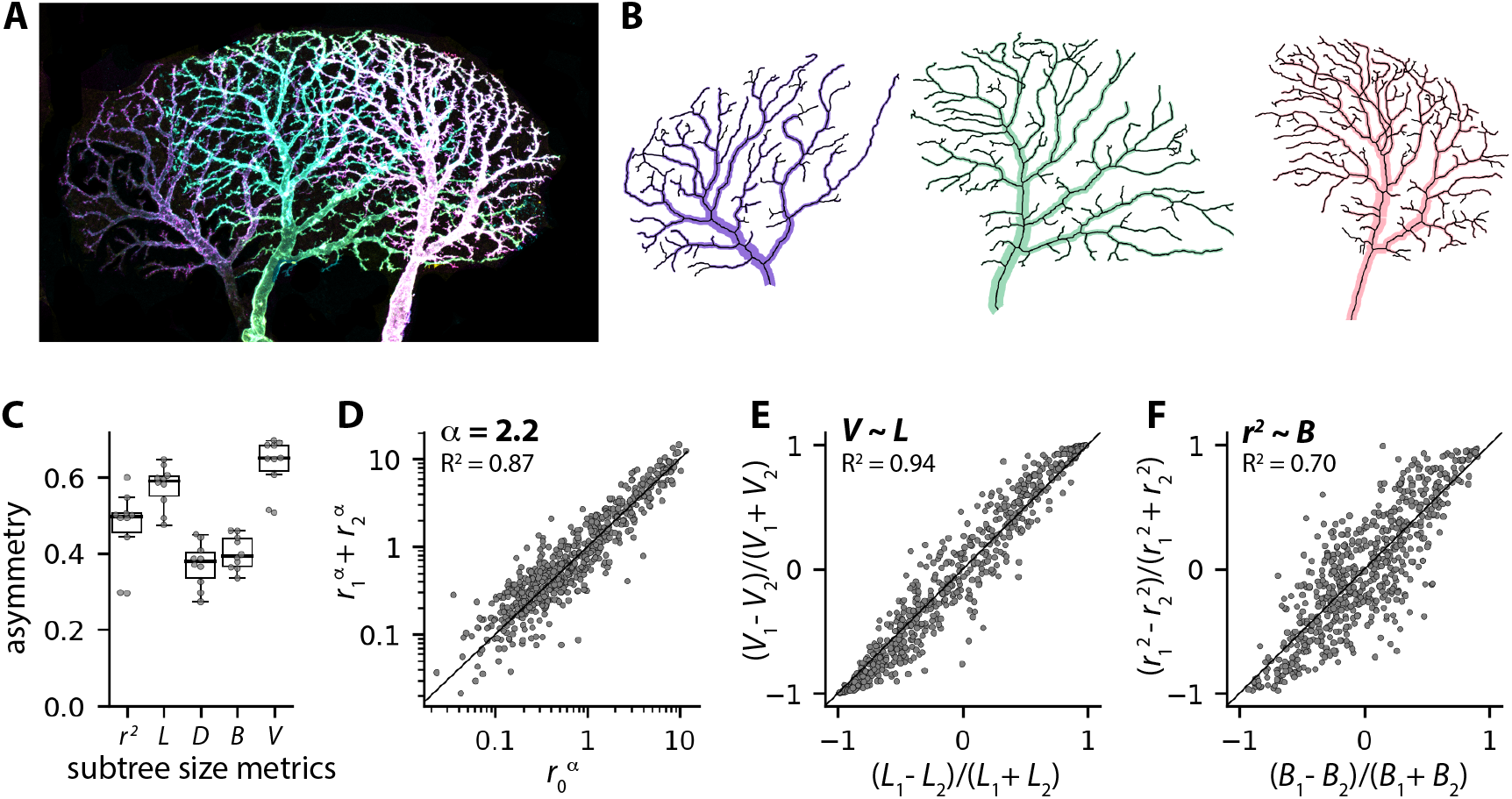
HS dendrites obey dendritic scaling rules. A-B: MCFO image of three HS dendrites (A) and extracted skeletons and radii (B). C: Box plots showing sister subtree asymmetries in trunk thickness (*r^2^*), length (*L*), depth (*D*), bushiness (*B*), and volume (*V*). Black lines, boxes, and whiskers indicate the median, interquartile range, and 1.5 times the interquartile range, respectively, and each gray dot indicates the average value for a single cell (N = 10 HS dendrites). D-F: HS dendrites follow parent-daughter scaling *r_0_*^α^ = *r_1_*^α^+*r_2_*^α^, with α = 2.2 (D), and sister subtree scaling with subtree volume proportional to subtree length (E) and trunk thickness (*r*^2^) proportional to subtree bushiness (F). N = 649 branch points from 10 dendrites.

Next, our model predicts that Da Vinci-scaled dendrites can only achieve equitable distribution of mitochondria if sister subtrees have volumes proportional to their length. To test this prediction, we compared the asymmetry in subtree lengths (*L_1_-L_2_*)/(*L_1_+L_2_*) with the asymmetry in volumes (*V_1_-V_2_*)/(*V_1_+V_2_*) for sister subtree pairs emerging from the same branch point. We found that length asymmetry is equal to volume asymmetry (Figure 6E, R^2^=0.94), indicating that longer sister subtrees have proportionally larger volumes, as predicted. Thus, HS dendrites obey two separate morphological rules: power law scaling of parent and daughter branches with α ~ 2, and sister subtree splitting with volume proportional to length (*L_1_/V_1_* = *L_2_*/*V_2_*). According to our model, for dendrites that follow these two rules, daughter branch cross-sectional areas must be proportional to subtree bushiness: *r_1_^2^/r_2_^2^* = (*L_1_*/*D_1_*)/(*L_2_*/*D_2_*). To test this prediction, we compared asymmetry in branch cross-sectional area (*r_1_^2^*-*r_2_^2^*)/(*r_1_^2^*+*r_2_^2^*) to asymmetry in subtree bushiness ((*L_1_*/*D_1_*)-(*L_2_*/*D_2_*))/((*L_1_*/*D_1_*)+(*L_2_*/*D_2_*)). We found that trunk cross-sectional area and bushiness asymmetry are correlated (Figure 6F, R^2^=0.70), and that cross-sectional area asymmetry was only weakly correlated with subtree length or depth asymmetry (Figure S6A-B). Finally, we measured similar correlations for synthetic tree structures with an imposed error in radii measurements (Supplemental Figure S6).

Altogether, these experimental results support our model and strongly suggest that dendrite morphology plays a key role in determining steady-state mitochondrial localization patterns in neurons *in vivo*.

### Robust self-organization of equitably distributed mitochondria for a range of transport parameters

In our model, we assume that mitochondrial arrest rates are inversely proportional to the cross-sectional area of each dendritic branch (*k_s_* ~ 1/*r*^β^, where β = 2), resulting in increased mitochondrial densities in thinner distal dendrites. However, distal enrichment was orders of magnitude higher in our model (~300 fold enrichment) compared to in our experimental measurements (~4 fold enrichment). Smaller values of β should result in more reasonable distal enrichment, so we estimated β from *in vivo* images of mitochondria moving through dendritic branches with a range of radii (see Methods). Our measurements show that mitochondrial arrest is indeed more frequent in thinner branches, and that the rate of arrest scales with dendrite radius according to *k_s_* ~ 1/*r^1.3^* (β = 1.3, R^2^ = 0.54, 95% confidence interval = 0.93-1.67) (Figure 7A). In our model, values for α and β in the range of our experimental measurements result in both realistic distal enrichment (~10 fold enrichment) and equitable mitochondrial densities in dendrites that obey r^2^~L/D sister subtree scaling (Figure 7B-C). Incorporating a minimum radius (r_m_) such that parent and daughter dendrites scale according to r_0_^2^+r_m_^2^ = r_1_^2^+r_2_^2^ (as recently described for sensory neurons in *Drosophila* larva) (Liao et al., 2021) reduces distal enrichment and introduces small asymmetries in mitochondrial densities across sister subtrees (Figure S7A-B). Interestingly, in Da Vinci-scaled dendrites (α = 2) without a minimum radius, equitable distribution is highly robust to changes in β (Figure 7C), indicating that changes in transport parameters have a dramatic effect on distal enrichment without affecting equitable distribution. In contrast, Rall’s (α = 3/2) and Murray’s (α = 3) laws failed to yield equitable mitochondrial distributions (Figure 7C), and different sister subtree scaling rules (e.g. trunks splitting according the subtree length, rather than bushiness) yielded equitable distributions in Da Vinci-scaled dendrites only when β = 0 (Figure S7D,F). Thus, equitable mitochondrial distributions are robust to changes in mitochondrial transport parameters only in dendrites that follow specific morphological scaling rules.

**Figure 7:**
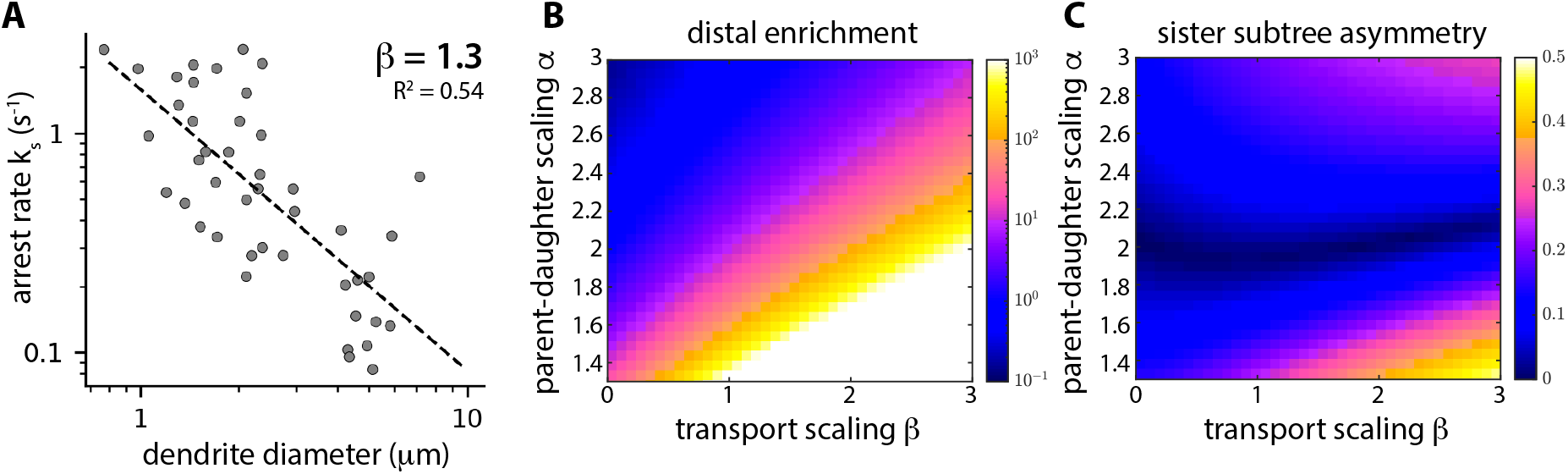
Equitable mitochondrial distributions are robust to changes in transport scaling. A: Dendrite diameters plotted versus mitochondrial arrest rate k_s_. The dashed line indicates the best fit, with k_s_ ~ 1/r^1.3^. B-C: Model results showing average distal mitochondrial enrichment (B) and mitochondrial density asymmetry across sister subtrees (C) calculated as a function of α (parent-daughter scaling) and β (transport scaling). HS dendrite topologies were extracted from MCFO images (N = 10 cells) and dendrite radii were set according to sister subtree scaling rule r^2^~L/D, as well as power law parent-daughter scaling with the indicated range of values for α.

## DISCUSSION

Neuronal function is inextricably linked to neuronal form: dendrite size limits the formation of synaptic connections to a particular receptive field (Peichl and Wassle, 1983); branching patterns affect dendritic integration of input signals (van Elburg and van Ooyen, 2010; Lefebvre et al., 2015); and axon thickness regulates action potential propagation (Waxman and Bennett, 1972; Seidl, 2014). Our work suggests that HS neurons in the *Drosophila* visual system also maintain a dendritic architecture that allows reliable distribution of mitochondria through asymmetrically branched arbors. We present a model in which four simple scaling rules determine steady-state mitochondrial distribution patterns (Figure 4). The first two rules — scaling of mitochondrial transport with dendrite radius and proportional splitting of mitochondria at branch points — relate local mitochondrial motility rates to dendritic branch radii. The third and fourth rules — power law scaling of parent and daughter radii and scaling of trunk thickness with sister subtree size — are morphological rules that determine the architecture of the dendrite. There are many possible forms of these dendritic scaling rules, but only a subset of the rules we examined — Da Vinci scaling of parent-daughter radii at branch points and sister subtree scaling with trunk thickness proportional to subtree bushiness — predict realistic mitochondrial localization patterns in our model. Our experimental measurements demonstrate that HS dendrites do in fact obey these morphological scaling rules (Figure 6). Thus, our work suggests that intracellular transport, and the need to distribute mitochondria throughout elaborately branched dendritic arbors, acts as an important constraint on dendrite morphology.

We have shown that mitochondria are equitably distributed across sister subtrees and enriched in the distal dendrites in HS cells (Figure 1). Distribution of mitochondria throughout the cell is critical for neuronal stability (Griparic et al., 2004; Verhoeven et al., 2006; Baloh et al., 2007; Nunnari and Suomalainen, 2012) but the relationship between specific mitochondrial localization patterns (e.g. distal enrichment) and neuronal function is unclear. One possibility is that mitochondrial densities simply reflect local energetic demands, with mitochondrial enrichment in subcellular regions that require relatively higher levels of ATP production. The reversal of ion fluxes near synapses is thought to account for a large fraction of the neuron’s energy budget (Attwell and Laughlin, 2001; Harris et al., 2012), and mitochondrial densities weakly correlate with synaptic densities in the dendrites of mouse cortical pyramidal neurons (Turner et al., 2022). Mitochondria also buffer calcium (Werth and Thayer, 1994), and variations in mitochondrial densities may contribute to compartment-dependent differences in calcium buffering capacities, which have recently been shown to contribute to place field formation in awake and behaving mice (O’Hare et al., 2022).

In addition to supporting dendritic function, enrichment of mitochondria in specific neuronal compartments may play a role in supporting mitochondrial function (Chang et al., 2006; Thomas et al., 2019; Turner et al., 2022). In active mitochondria, damaging reactive oxygen species are a by-product of the electron transport chain (Miwa and Brand, 2003). Mitochondria are thought to compensate for ROS-induced damage in two ways: by degrading and replacing damaged proteins (Wang et al., 2012), and by homogenizing the mitochondrial population — diluting damaged proteins and sharing freshly synthesized proteins — via mitochondrial fusion and fission (Willems et al., 2015). The majority of mitochondrial proteins are thought to be synthesized and transported into mitochondria in the cell body which are then trafficked out into the axons and dendrites (Misgeld and Schwarz, 2017). Theoretical work suggests that fusion with stationary mitochondria depletes freshly-synthesized mitochondrial proteins from motile mitochondria as they move in the anterograde direction (Agrawal and Koslover, 2021). A graded distribution of stationary mitochondria, with higher densities in distal dendrites, may allow neurons to ensure adequate delivery of fresh mitochondrial proteins to distal axons and dendrites while also maximizing complementation across mitochondria in distal compartments. Moreover, mitochondria in HS distal dendrites are large, often spanning multiple dendritic branch points (Figure S1). If young, healthy mitochondria fuse with stationary mitochondria upon arrival in the distal dendrites, passive transport within these elongated mitochondria would ensure uniform local distributions of freshly-synthesized mitochondrial proteins. Future versions of our model will include mitochondrial fusion and fission rates, as well as mitochondrial motility.

In our model, equitable distribution of mitochondria across sister subtrees is robust to variation in mitochondrial transport parameters in Da Vinci-scaled dendrites (Figures 7C). Distal enrichment, on the other hand, depends on inverse scaling of motility arrest with dendrite thickness (Figure 7B). The mechanism underlying inverse scaling of motility arrest and branch thickness remains undetermined. In principle, narrowing of dendrite branches, on its own, could be sufficient to increase motility arrest. In cylindrical dendrites, the surface area-to-volume ratio (A_s_/V) increases as radius decreases, with A_s_/V ~ 1/r. Microtubule densities are conserved throughout dendritic arbors (Hillman, 1979; Kubota et al., 2011; Katrukha et al., 2021) and the amount of microtubule-based transport should scale with dendrite volume. In contrast, some mechanical interactions that oppose mitochondrial transport should scale with surface area. Non-specific viscous friction between motile mitochondria and the cell membrane could oppose motility in thin neuronal processes (Narayanareddy et al., 2014). Actin localizes to the cell membrane in neurons (Leiss et al., 2009), and actin-based anchoring opposes microtubule-based transport in several contexts (Kapitein et al., 2013; Lu et al., 2020), including myosin V-dependent opposition to mitochondrial movement in cultured neurons (Pathak et al., 2010). Biochemical signals generated at the cell membrane could also contribute to inverse scaling of motility arrest and dendrite radius. For example, high glucose levels trigger mitochondrial arrest in cultured neurons via post-translational modification of the Milton adaptor protein (Pekkurnaz et al., 2014), and quantitative modeling indicates glucose-mediated motility should be sufficient to affect mitochondrial localization patterns (Agrawal and Koslover, 2021). Neurons take up glucose via transporters in the cell membrane (Ferreira et al., 2011; Lundgaard et al., 2015; Ashrafi et al., 2017) and, assuming a constant areal density of these transporters, glucose concentrations in the cytosol will increase as the surface area-to-volume ratio increases, thereby promoting increased mitochondrial arrest in thin distal dendrites. Altogether, the relative weight of mechanical and biochemical signals generated at the cell membrane versus in the cytosol should increase as neuronal processes narrow, and surface area-to-volume ratios may play a general role in regulating intracellular transport in neurons.

Finally, our results suggest that neuronal signal processing and housekeeping requirements may act as competing constraints on neuronal architecture. Specifically, HS dendrites are approximately Da Vinci-scaled (α ~ 2) which, according to our model, allows for equitable distribution of mitochondria across sister subtrees for a broad range of transport parameters. However, according to cable theory, parent-daughter scaling according to Rall’s law (α = 3/2) is optimal for dendritic function, as it allows for efficient propagation of electrical signals across branch points in passive dendrites (Rall, 1959). Interestingly, unlike many neurons in the *Drosophila* visual system, HS neurons are not purely graded (Schnell et al., 2010), and active dendritic conductances may allow Da Vinci-scaled dendrites to efficiently integrate dendritic inputs while also maintaining steady-state mitochondrial localization patterns. Parent-daughter scaling may be more likely to be constrained by Rall’s law in neurons with purely passive dendrites. Strikingly, though, our model strongly suggests that Rall’s law is incompatible with equitable distribution of mitochondria in asymmetrically branched dendrites; for equitable mitochondrial distribution, Rall-scaled dendrites must also be symmetrically branched. Altogether, our work suggests that neuronal morphologies are constrained not only by functional requirements and wiring economy, but also by the need to efficiently distribute subcellular constituent elements like mitochondria throughout the cell.

## METHODS

### *Drosophila* strains and husbandry

The *Drosophila* stocks used in this study are listed in the Key Resources table. All flies were reared at 25°C on standard cornmeal-agar food in 12 h light: dark cycle. Crosses were flipped into fresh vials every 3 days and progeny were imaged 4-7 days after eclosion.

### *In vivo* imaging

Female flies were cold anesthetized and positioned in a key-hole cut in a thin metal shim, with the back of the head exposed above the shim and the eyes below the shim. The fly was secured in place with fast-curing glue (Bondic) and the brain was exposed using fine forceps to dissect a hole in the cuticle and remove overlying fat and trachea. The brain was perfused with a sugar saline solution (103 mM NaCl, 3 mM KCl, 5 mM TES, 1 mM NaH_2_PO_4_, 26 mM NaHCO_2_, 4 mM MgCl_2_, 1.5 mM CaCl_2_, 10 mM trehalose, 10 mM glucose, and 7 mM sucrose). Neurons were imaged using an integrated confocal and two-photon microscope (Leica SP8 CSU MP Dual) and a 25x 1.0 NA water immersion objective (Leica). For confocal imaging of motile mitochondria, stationary GFP-tagged mitochondria in the field of view were photobleached prior to time lapse imaging, allowing for resolution of individual motile mitochondria as they moved into the field of view. Confocal z-stacks (voxel size = 108.54 nm x 108.54 nm x 22 microns) were collected every 4.4 seconds for 10-20 minutes after photobleaching. For two-photon imaging of visual stimulus-evoked calcium signals, a fixed 1045nm femtosecond laser beam (Spectra-Physics Insight X3 DUAL) was used to excite RGECO1a, and 256 × 256 images (pixel size = 139.09nm x 139.09nm) were collected at a frame rate of 10 Hz.

### Visual stimulus presentation

Visual stimuli were generated using PsychoPy (Python) and presented on a white screen (Da-Lite Dual-Vision vinyl, AV Outlet) using a digital light projector (DLP LightCrafter, Texas Instruments). The stimulus screen spanned ~60° of the fly’s visual field horizontally and ~60° vertically, and the stimulus was updated at 60 Hz. To avoid detection of light from the stimulus by the microscope, the stimulus was filtered using a 472/30 nm bandpass filter (Semrock). Voltage signals from the imaging software were relayed to PsychoPy via a LabJack device, in order to synchronize the stimulus and the imaging frames. The visual stimuli were full contrast square wave gratings (Λ = 30°) that filled the entire stimulus screen. When the stimulus was on, the gratings moved in the preferred direction for HS neurons (front-to-back across one eye) at 30°/s; when the stimulus was off, the gratings remained stationary.

### MultiColor FlpOut Labeling

HS Gal4 driver lines were crossed with MCFO virgins. Offspring were collected 1-2 days after eclosion, heat shocked at 38°C for 25 min, and dissected three days later. Fly brains were dissected in cold PBS solution and fixed in 4% formaldehyde for 25 min at room temperature. Brains were subsequently rinsed with PBST (PBS with 0.5% Triton) and blocked in PBST-NGS (PBST with 5% normal goat serum) at room temperature for 1.5 hr. Brains were incubated for two nights in primary antibodies diluted in PBST-NGS, then incubated for two nights in secondary antibodies in PBST-NGS, and finally incubated overnight in tertiary antibodies in PBST-NGS. Prior to each antibody incubation, brains were washed 3 times for 10 min each in PBST. All antibody incubations were performed at 4°C. Brains were mounted on their ventral end in VectaShield media (Vector Laboratories) and imaged using confocal microscopy. All antibodies are listed in the Key Resources Table.

### Quantification of mitochondrial morphologies and localization patterns

Mitochondrial morphologies and distribution patterns in HS neurons were measured using the publicly available fully aligned fly brain (FAFB) dataset, serial section transmission electron microscopy (ssTEM) images of the entire brain of an adult female fly (Zheng et al., 2018). Previously traced HS skeletons (Michael Reiser, unpublished data), available on the CATMAID (Collaborative Annotation Toolkit for Massive Amounts of Image Data) server (Saalfeld et al., 2009), were used to identify HS neurons within the larger FAFB image volume. To measure the size of individual mitochondria, small image volumes centered around HS dendritic segments were cropped out of the FAFB dataset, and mitochondria within HS dendrites were manually segmented in three dimensions using the TrakEM Fiji plug-in. To measure mitochondrial localization patterns throughout HS neurons, all HS skeletons were resampled using a python-CATMAID interface library, pymaid, such that the graph distance between skeleton nodes was 5 microns. All branch points and end points were preserved during resampling. Two dimensional image slices centered around each node in the resampled skeleton were then cropped out of the FAFB dataset, and HS neurons and the mitochondria within them were manually segmented in each image using TrakEM. Mitochondrial density (total mitochondrial area/total neurite area) was measured as a function of neuronal compartment (axons versus dendrites), distance from the soma, and across sister subtree pairs.

### Quantification of mitochondrial motility patterns

Mitochondrial speeds, flux rates, arrest rates, and lengths were measured from max projections of confocal z-stacks of mitoGFP and cytosolic tdTomato expressed in HS neurons. Max projections were aligned using the Turboreg Fiji plugin. Individual motile mitochondria were hand-tracked using the Tracking Fiji plugin, and mitochondrial speeds were measured from mitochondrial tracks using custom-written Python code. Linear mitochondrial flux rates were measured by counting the number of motile mitochondria that moved through a particular cross-section of a dendritic branch in either the anterograde or retrograde direction per unit time. Mitochondrial arrest rates were calculated per dendritic branch, as the fraction of mitochondria that either stopped or paused motility within each branch. Lengths of motile mitochondria were measured using Celltool (Pincus and Theriott, 2007), and the average volume of motile mitochondria was estimated as V = l pi r^2, where l is ~2 microns, measured from our *in vivo* images, and r is assumed to be ~0.3 microns. Dendrite diameters were measured based on the cytosolic tdTomato signal, using the line scan tool in Fiji.

### Quantification of dendritic branching parameters

Dendritic arbors for individual HS neurons were segmented from MCFO images using ilastik (Berg et al., 2019), publicly available interactive machine learning software, and custom-written Python code. Unique pixel classifiers were trained in ilastik for each MCFO z-stack, and binary masks were generated from the resulting probability maps in Python by using thresholding and connected component analysis. Binary masks were then manually cleaned up in Fiji and skeletonized using the Skeletonize (2D/3D) Fiji built in plugin.

Skeleton data was translated into a set of nodes (including junction nodes, parent node, and distal tips) with three-dimensional coordinates, and curved edge paths connecting the nodes. Once the initial network structure was extracted, manual clean-up was carried out with a custom Matlab GUI, involving the removal of short spurious branches (‘shrubs’) from the network. A combination of percentile and asymmetry cutoffs were used to quantitatively remove shrubs that would not contribute to the total length in subsequent analysis. A degree of manual editing was performed for each cell, such that any branch without a discernible thickness was removed from the network object. The widths of network branches were also calculated with the aid of the Matlab GUI. The interface allows the user to add and adjust width measurements across a given edge. For longer edges, one measurement point close to the branching point and the other closer to the end of the edge are chosen. Total subtree length, volume, depth, and bushiness following each branch was calculated using Matlab written code. Volume was measured using the diameter and length of each edge bounded between two nodes. Bushiness is defined as the total subtree length (L) over the subtree depth (D), where D is the path length from the base of the subtree to each distal tip, weighted by the length of each subtree. Lastly, asymmetry for each parameter was measured using our asymmetry metric, (ST1-ST2)^2^/(ST1+ST2)^2^, where ST1 and ST2 are the indicated measurements (radius, length, volume, or bushiness) for sister subtree 1 and 2, respectively.

### Synthetic tree construction

Synthetic binary trees were constructed in Python 3.7.6 using the NetworkX library (Aric A. Hagberg, 2008). First, the skeleton of a binary tree was created, starting with a single junction consisting of a parent branch and two daughter branches of unit length. Moving downstream along the tree, each daughter branch either terminated as a distal tip (with probability ⅓), increased in length by an additional unit (probability ⅓), or branched into two more daughter branches (probability ⅓). This process was repeated up to a preset maximum path distance (40 unit branch lengths) from the arbor parent node to the distal tips. The resulting random-topology binary tree structures were used as the ensemble of synthetic arbors in Supplemental Figures S5 and S6.

### Computing imposed radii

Skeletons with well-defined branch lengths and connectivity were obtained either from MCFO images of *Drosophila* HS neurons, from published data (Cuntz et al., 2013), from HS skeletons traced through a ssTEM dataset (Zheng et al., 2018) (Michael Reiser lab unpublished data or from synthetically constructed trees.

For each skeleton, the widths of the branches (r_i_) were calculated, starting with r_0_=1 at the parent trunk (in dimensionless units). At each junction node, the daughter branch radii were defined by a combination of scaling rule (1) for parent and daughter radii: r_1_^α^ + r_2_^α^ = r_0_^α^ and rule (4) for sister subtree radii r_1_^2^/ r_2_^2^ = μ_12_. Da Vinci arbors have α, = 2, Rall’s Law arbors have α, = 3/2, and Murray’s Law arbors have α, = 3.

The daughter branch splitting rules included equal splitting (μ_12_ = 1), splitting in proportion to total subtree length (μ_12_ = Σ_i∈ST1_ l_i_ / Σ_i∈ST2_ l_i_), and splitting in proportion to subtree bushiness (total branch length over depth), defined in Supplemental Methods 1.

To impose radii with a minimum radius (Liao et al., 2021) on an MCFO HS skeleton (Figure S7), we assumed the primary dendrite radius = 3 μm and the minimum radius Rm = 0.2 um (Liao et al., 2021). At each branching point, the parent-daughter radii relationship was modified to r0^2 + rm^2 = r1^2 + r2^2 to incorporate the effect of this minimum radius. A ‘minimum area’ is imposed to take into account the finite area occupied by a microtubule, so the distal dendrites have a greater predicted diameter in this case.

To estimate the effect of measurement error in branch width (Supplemental Figure S6), we added a Gaussian noise term with standard deviation equal to 30% of the computed radius to all branch widths ri.

### Mean-field model for mitochondrial distribution

Our minimal mean-field model allows the calculation of steady-state mitochondrial densities along each branch of a dendritic arbor with prescribed topology, branch lengths, and branch radii. We assume punctate mitochondria are produced at the soma (parent node of the arbor), and undergo processive bidirectional motion with pause free velocity v_i_ (where i is the branch index), pausing rate k_i_^s^, and constant restarting rate k_w_. The linear densities of anterograde, retrograde, and stationary mitochondria in each branch are given by ρ_i_^+^, ρ_i_^−^, ρ_i_^s^, respectively. At steady-state, these densities obey the transport equations:

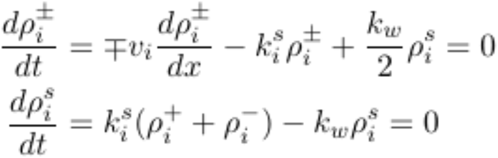

The steady-state solutions have densities that are constant along each individual branch, with the relationship between different branch densities determined by boundary conditions at the junctions. Distal tips are treated as reflecting boundaries (yielding ρ_i_^+^= ρ_i_^−^). At the branch junctions, the boundary conditions are set by the conservation of incoming and outgoing mitochondrial flux, together with the assumption that anterograde mitochondria split in proportion to the cross sectional area of the daughter branches:

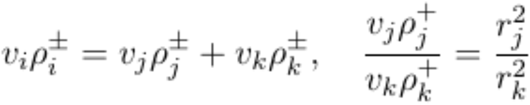

where i is the parent branch, and j, k the two daughter branches at the junction. Finally, we set the boundary condition at the soma by fixing the motile mitochondria linear density in the parent trunk to a constant, ρ_0_.

The steady-state linear densities of mitochondria in each branch are then found by solving this set of linear equations. The density of stationary mitochondria is given by:

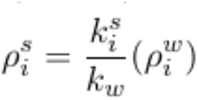

where ρ_i_^w^= ρ_i_^+^ + ρ_i_^−^ is the motile mitochondrial density.

### Volume densities and equitability metric

Volume density on each branch is computed as c_i_ = ρ_i_/r_i_^2^. The average volume density in a subtree is given by:

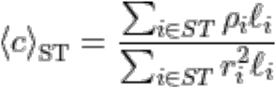

Where the summation is carried out over all branches i of the subtree, with corresponding length l_i_ and radius r_i_.

We define a single metric ζ for the equitability of mitochondrial distribution throughout the entire tree. Specifically, this metric gives the root-mean-squared asymmetry of mitochondrial densities in sister subtrees, averaged over all junctions in the arbor. We focus on the regime where most mitochondria are in the stationary state, and hence compute the asymmetry metric based specifically on distributions of stationary mitochondria:

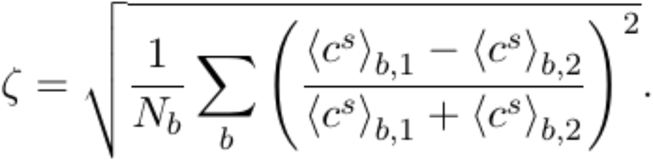

Here, the index b enumerates the junctions, N_b_ is the total number of junctions in the tree, and subscripts 1 and 2 refer to the two daughter subtrees emanating from a junction.

### Agent-based simulations of mitochondrial transport

Stochastic simulations of mitochondrial transport in a network were carried out using custom code written in Fortran 90. An initial tree structure is initialized to contain information on the junction connectivity, as well as lengths and radii of individual branches. The tree skeletons were generated using the NetworkX library as described in (Synthetic tree construction). A typical example is presented in Supplemental Figure S5B. The widths of these synthetic trees were calculated by imposing either Da-Vinci scaling law with r^2^ ~ *L/D*, or Rall’s law with *L ~V* scaling applied. For demonstrating the simulated results showing the difference in equitability of mitochondrial distribution in trees following Da-Vinci vs Rall’s scaling, *N = 10* synthetic trees were used. The edge lengths for the synthetic tree were fixed to *l_0_ = 10 μm*, consistent with average edge length measurements observed in HS dendrites.

While these synthetic trees were shorter in extent compared to real HS neurons due to computational limitations (going a maximum of *D/l_0_ = 4* levels down), the synthetic trees incorporated a range of heterogeneity in branching structure.

The tree is then populated by a fixed number (*N = 1000*) of punctate of mitochondria, distributed uniformly throughout. The position of each mitochondrion is tracked in terms of the branch on which it is located and its position along the branch. Each mitochondrion is associated with a transport state (anterograde, retrograde, or stationary).

On every timestep, an anterograde mitochondrion steps distance *vΔt* downstream along the branch, and a retrograde mitochondrion steps distance *vΔt* upstream. The velocity of an individual mitochondrion was assumed to be *v = 0.4 μm/s*. One time step corresponds to *Δt = 2.5 s*, so that the stepping distance *vΔt = 1 μm*. A motile mitochondrion switches to a stationary state with probability P_stop_, whereas a stationary mitochondrion becomes motile with probability P_start_, according to the corresponding rates:

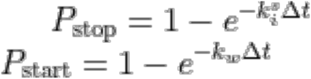

When becoming motile, the mitochondrion is equally likely to enter the anterograde or retrograde state. The rate of switching from motile to stationary state was fixed according to measurements of mitochondrial motility in HS dendrites. From the measurements of mitochondrial densities and exchange rate, we obtain a ~ 3% motile fraction in the primary dendrite. Plugging in to the expression for motile fraction *f_motile_* = 1 + *k_s_/k_w_*, we obtain the dimensionless ratio *k_s_/k_w_* = 35 for the primary dendrite. The rate *k_s_* for a primary dendrite was set equal to 0.15/s and the rate *k_w_* was set equal to 2 × 10^−3^/s, so that on an average a mitochondrion had a probability *P_stop_ =* 0.31 of stopping in a primary dendrite at each timestep. The stopping to walking rate was tuned as a function of branch radii according to the rule *k_s_* ~ 1/r^2^.

At each junction, an anterograde mitochondrion chooses which daughter branch to enter with probability proportional to the cross-sectional area of the branch: p_1_/p_2_ = r_1_^2^/r_2_^2^. When an anterograde mitochondrion reaches the distal tip, it reverses and becomes retrograde. When a retrograde mitochondrion reaches the soma, it becomes anterograde again. The simulations were run for *N_steps_ = 10^8^* to ensure convergence in observed mitochondrial distributions, as monitored by the edge density autocorrelation function over time.

### Estimating the power law rule for dependence of motility arrest on dendritic branch thickness

Figure 7A shows the dependence of the pausing rate *k_s_* on dendritic diameter, from experimental mitochondrial motility data. Each field in the motility data gave the number of transport events observed for a specific dendritic branch in the entire imaging duration, classified as (i) anterograde-persistent (ii) anterograde-stop (iii) anterograde-pause (iv) anterograde-reverse (v) retrograde-persistent (vi) retrograde-stop (vii) retrograde-pause and (viii) retrograde-reverse. The probability of stopping (*P_stop,i_*) for the branch *i* was calculated as the fraction of total stopping events (antero-stop + retro-stop + antero-pause + retro-pause) and total transport events recorded. From the probability of stopping, the effective stopping rate *k_s, i_* for each branch *i* was calculated from the formula *P_stop,i_* = 1 – *e^−k_s,i_Δt^*, where *Δt* is the time spent by mitochondria in that branch, which is proportional to the length of the branch.

---

**Figure S1:**
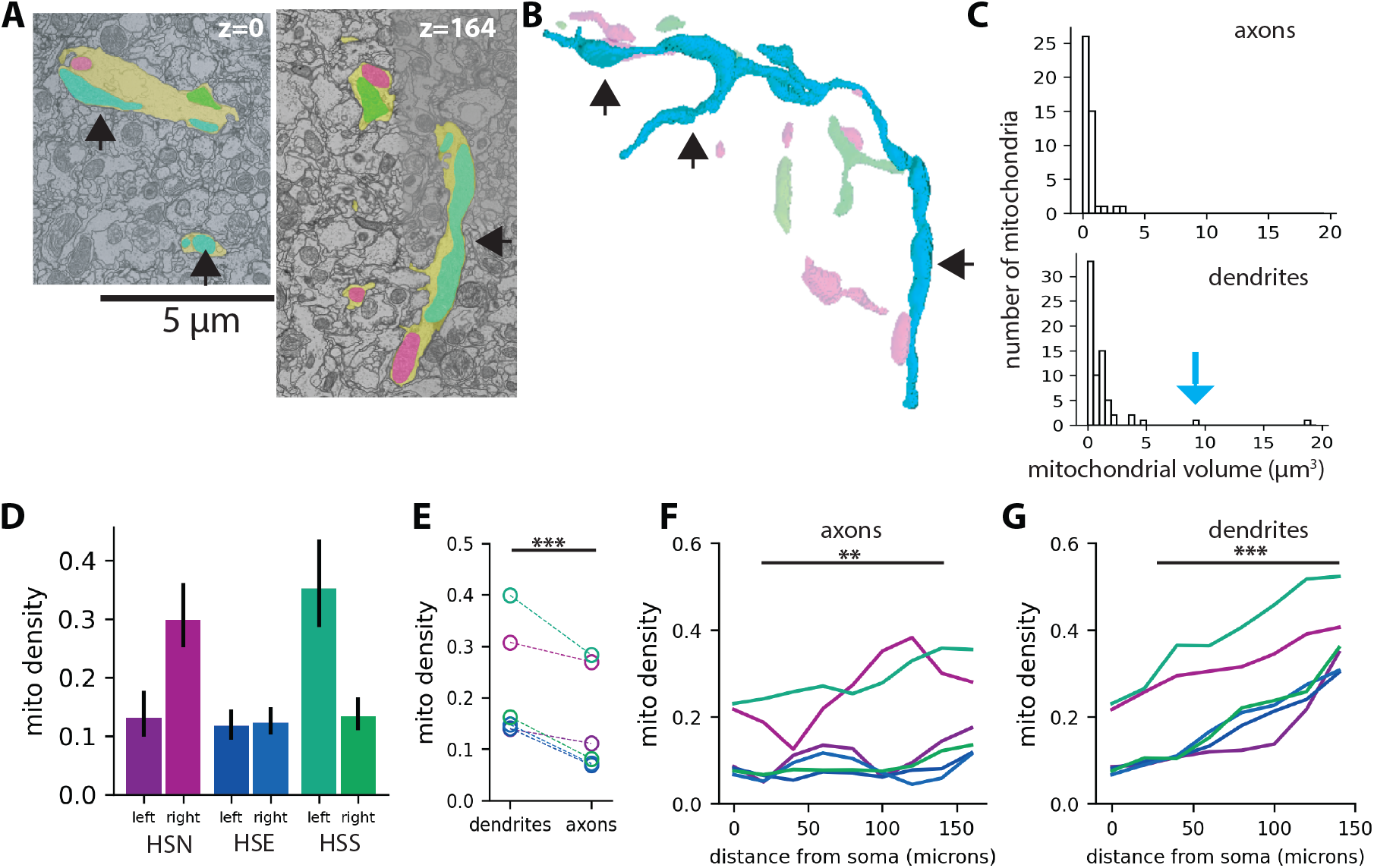
Mitochondrial morphologies and localization patterns in HS neurons. A: TEM images showing mitochondria (green, cyan, magenta) in an HS dendrite (yellow). B: 3D reconstruction of mitochondria based on ssTEM images. Black arrowheads point to different portions of a single branched mitochondrion. C: Mitochondrial volumes in axons (top, N = 53 mitochondria from 5 cells) and dendrites (bottom, N = 48 mitochondria from 5 cells). The blue arrow indicates the volume of the large mitochondrion shown in B. Dashed lines indicate median mitochondrial volumes (0.40μm^3^ in axons; 0.54μm^3^ in dendrites). D: Total mitochondrial densities for six HS neurons; error bars are 95% bootstrap confidence intervals. E: Mitochondrial densities in axons versus dendrites. F-G: Mitochondrial density plotted versus distance from the cell soma in axons (F) and dendrites (G). Colors indicate HS subclasses (magenta = HSN; blue =HSE; and green = HSS). Asterisks indicate significant differences (paired t-test; **p < 0.01 and ***p < 0.001).

**Figure S2:**
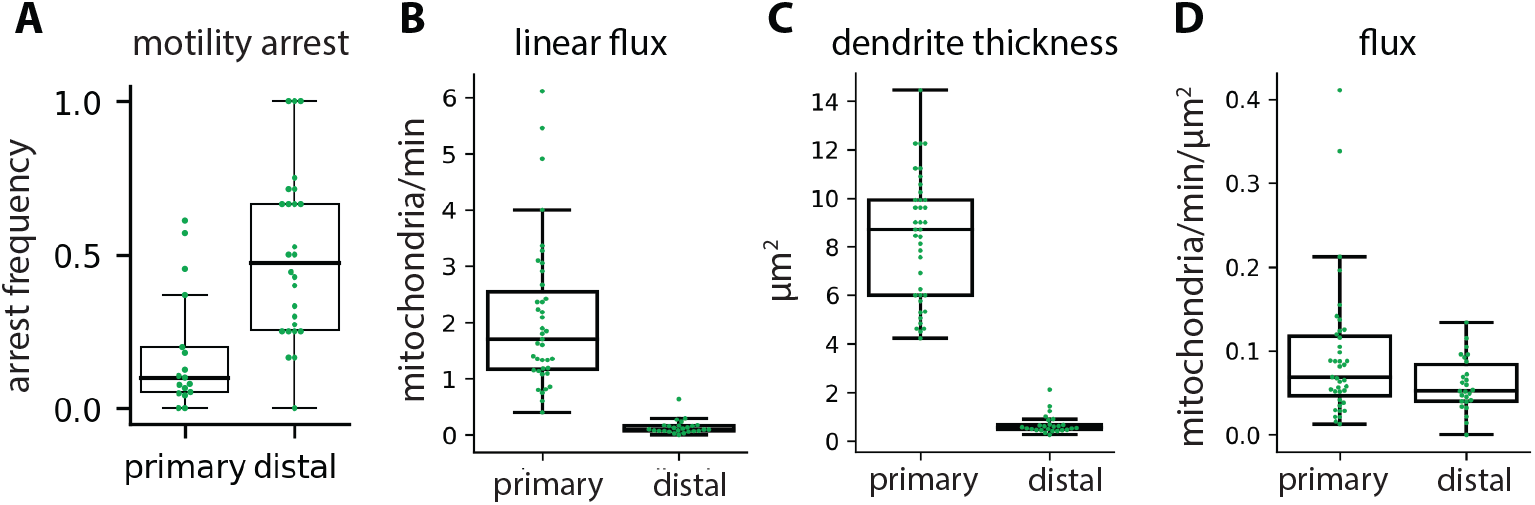
Anterogrde mitochondrial motility in primary and distal HS dendrites. Mitochondrial arrest rates (A), linear flux (B), dendrite thickness (C), and flux (D) in primary versus distal HS dendrites. Mitochondrial arrest rate is the fraction of mitochondria that arrested motility per branch. Each dot represents average metric per fly, N = 39 (primary dendrites) or 26 (distal dendrites) flies.

**Figure S3:**
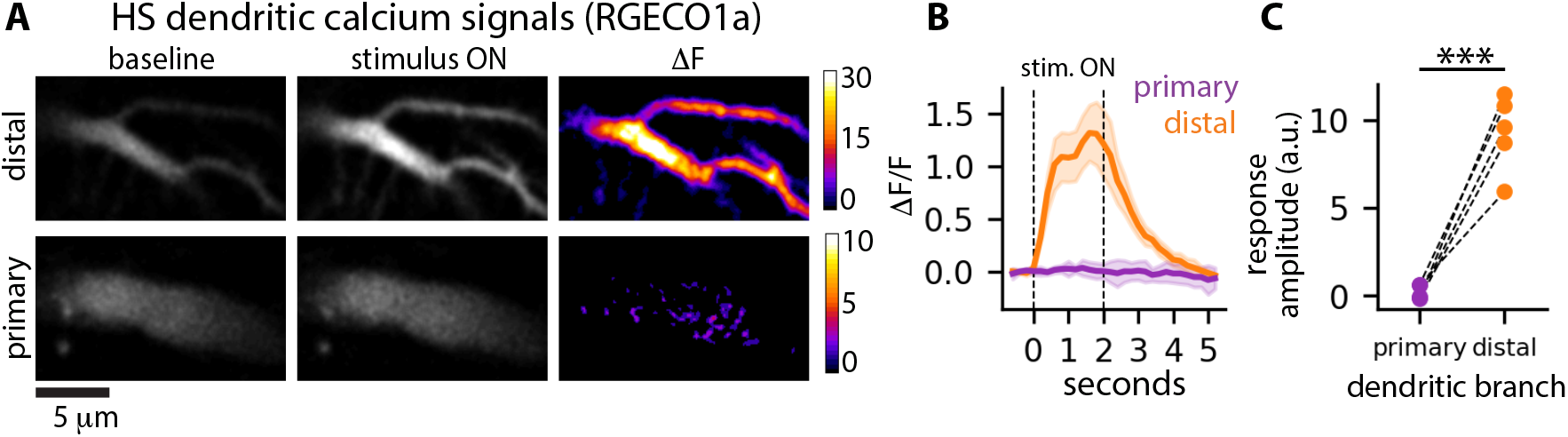
Visual stimulus-evoked calcium responses in HS primary and distal dendrites. A: Two-photon microscopy images of calcium (RGECO1a) signals in HS distal (top row) and primary (bottom row) dendrites before (baseline, left column) and during (stimulus ON, center column) visual stimulus presentation. The images on the right show the difference between the left and center images (ΔF = stimulus ON - baseline). The visual stimulus was a square wave grating moving in the preferred direction for HS neurons. B: RGECO1a responses to visual stimulation in primary (purple) and distal (orange) dendrites, plotted over time. The dashed lines indicate when the stimulus was on. C: Stimulus-evoked RGECO1a response amplitudes in primary and distal dendrites. Individual dots indicate average response amplitudes in one fly; dashed lines connect measurements in the primary and distal dendrites in the same fly. Asterisks indicte a significant difference (p<0.001, paired T test).

**Figure S4:**
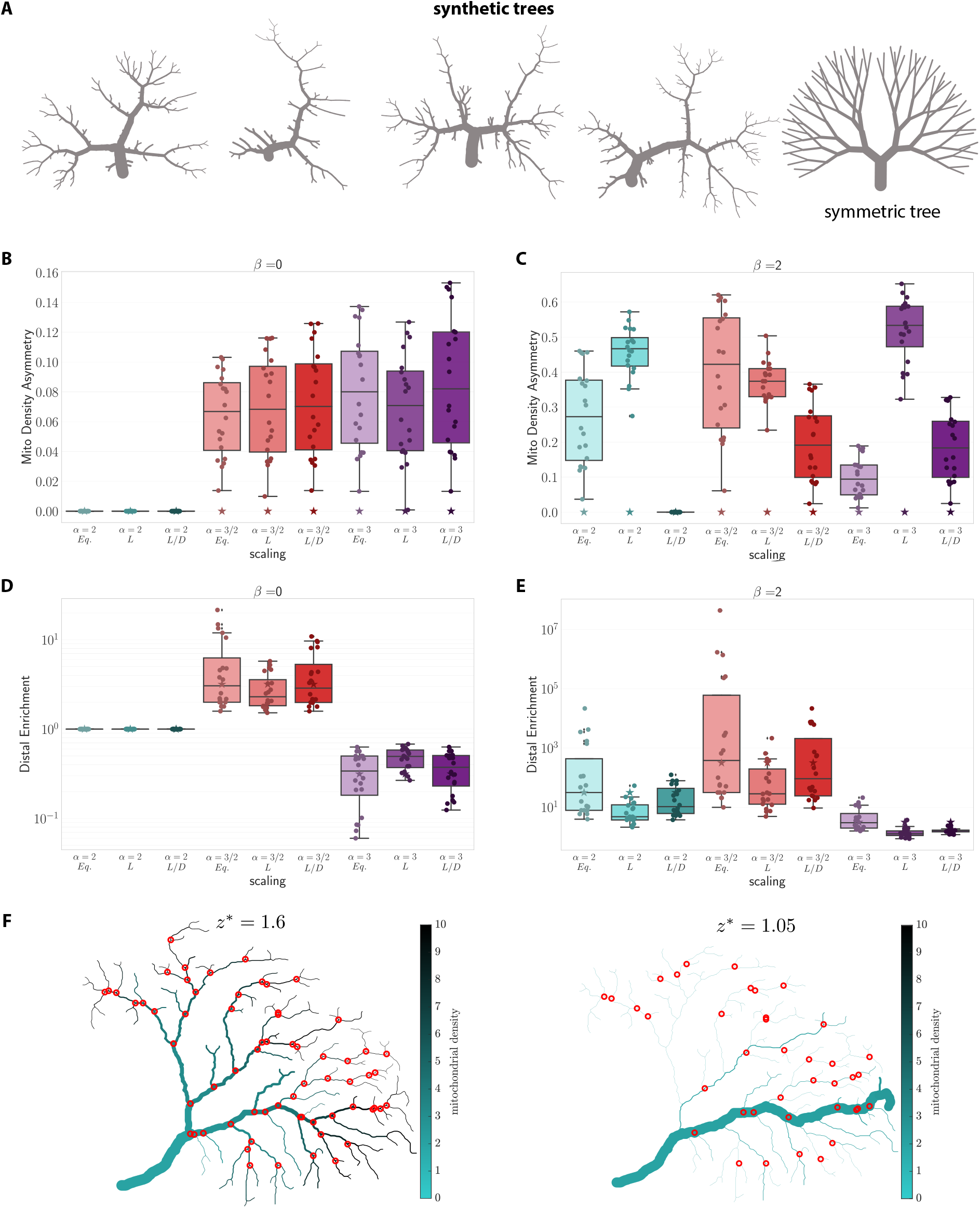
Model mitochondrial distributions in synthetic dendritic trees. A: Representative examples of synthetic trees with random (four left images) or symmetric (right) topologies. BE: Model results. Mitochondrial density asymmetry across sister subtrees (B,C) and distal mitochondrial enrichment (D,E) were calculated for β = 0 (B,D) or β = 2 (C,E), a = 2 (cyan), 3/2 (red), or 3 (purple), and sister subtree scaling with subtree trunks splitting according to r_1_=r_2_ (eq.), r^2^~L, or r^2^~L/D. N = 19 synthetic arbors (circles) and 1 symmetric arbor (stars). F: HS dendrites with radii that obey Rall’s parent-daughter scaling (a = 3/2); β = 0. The parameter z* determines the ratio of sister subtree trunk thickness (r_1_ and r_2_) at each branch point (see Supplemental Methods). Red circles indicate branch points for which there is no solution for equal sister subtree densities. For high values of z* (left), there are many junctions with no solution. As z* approaches 1 (right), there are fewer junctions with no solution. However, arbor morphologies become unrealistic, with many thin dendrites branching from a single thick branch.

**Figure S5:**
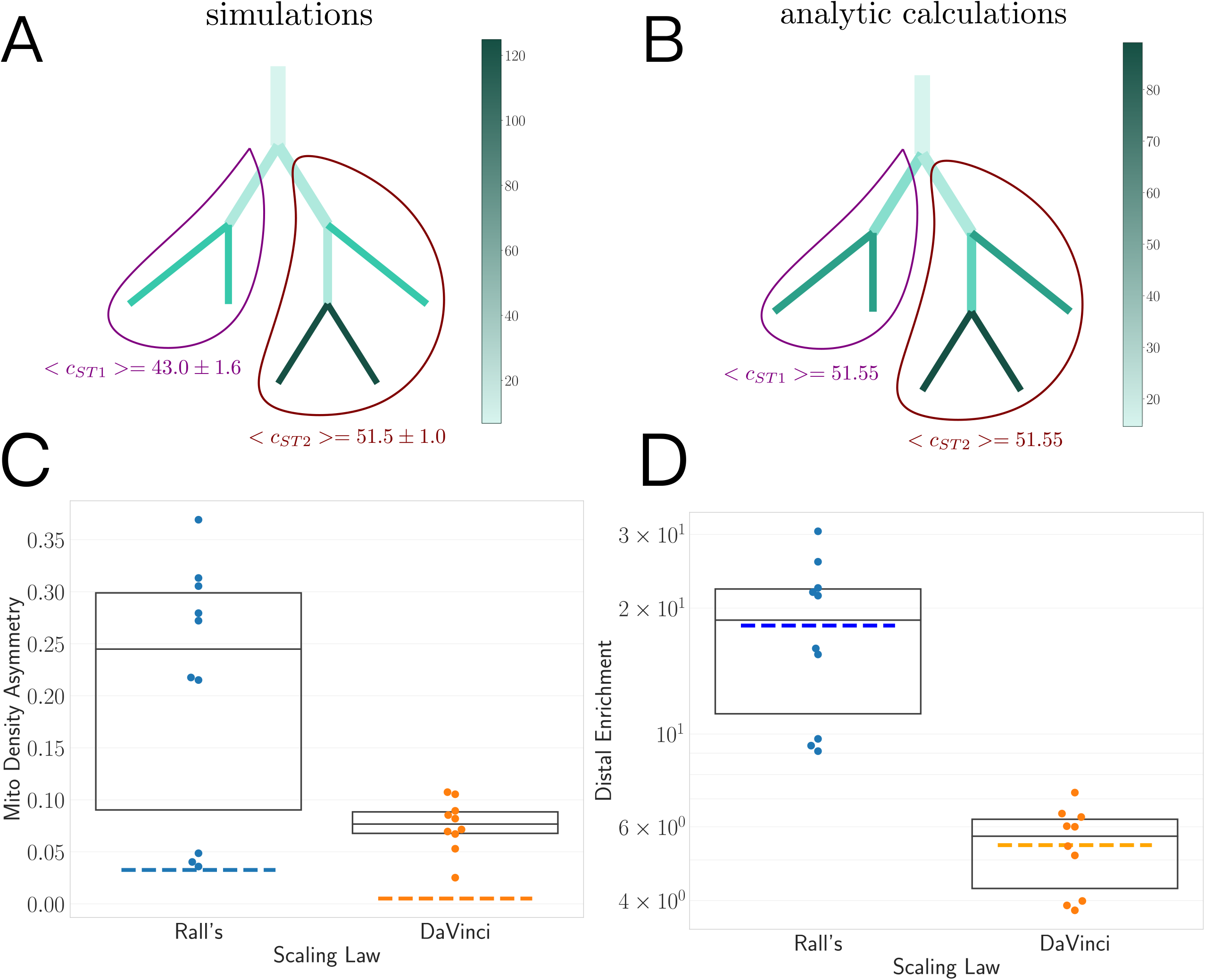
Numerical simulations of mitochondrial distributions. A: Simulated mitochondrial densities in a small synthetic arbor scaled according to Da Vinci’s rule (α = 2), sister subtree splitting with L~V, and transport scaling with β = 2. Total mitochondrial densities in the two largest subtrees (<c_ST1_> and <c_ST2_>) are indicated on the plot. B: Analytical calculations of mito-chondrial densities for the same arbor as in A. Note that whereas mitochondria are equitably distributed across ST1 and ST2 in the analytical solution, stochastic effects result in asymmetry in the simulated results. C-D: Mitochondrial densities were simulated for 10 synthetic arbors, and mitochondrial density asymmetry (C) and distal enrichment (D) were measured for two parent-daughter scaling rules: Rall’s law (〈 = 3/2) and Da Vinci’s rule (〈 = 2). Sister subtrees split with L~V, and transport scaled with β = 2. Box plots indicate the mean and interquartile range for simulation results; dashed lines indicate average values for numerical solutions for the same arbors.

**Figure S6:**
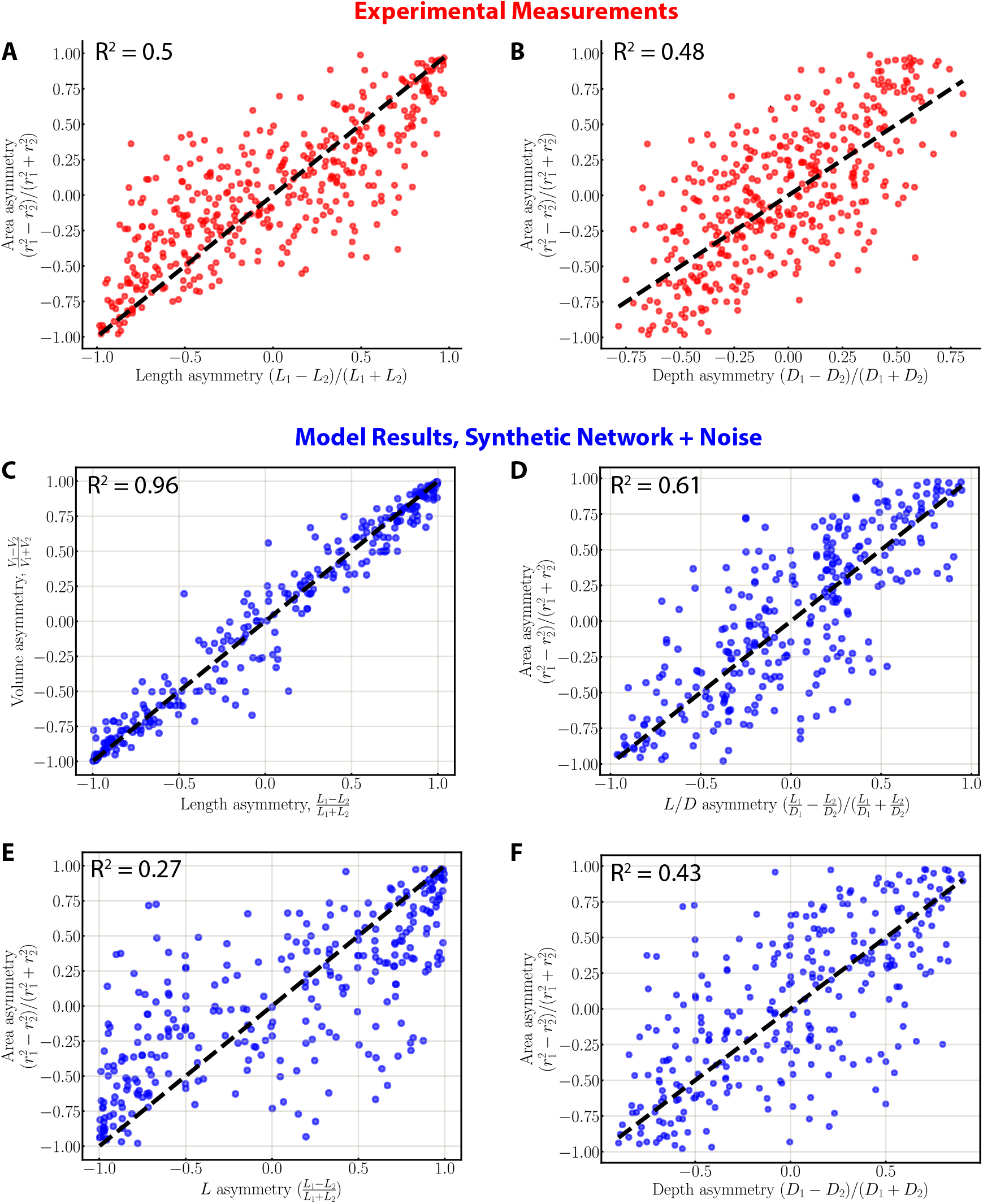
HS dendrites do not obey sister subtree scaling with r^2^ ~ subtree length or depth. A-B: Measurements of HS dendrites sister subtree asymmetries (N = 649 branch points from 10 HS dendrites). Trunk thickness (r^2^) asymmetry is more weakly correlated with length (A) or depth (B) asymmetry than with bushiness asymmetry (Figure 6F). C-F: Sister subtree correlations in a synthetic tree. Branch radii were set according to parent-daughter scaling with α = 2 and sister subtree scaling with L~V, plus a gaussian noise term (see Methods). The noise has a small effect on measurements of L~V scaling (C), and a larger effect on r^2^~L/D scaling (D). Subtree length (E) and depth (F) asymmetries show weaker correlations with r^2^ asymmetry, as in the experimental results (A-B).

**Figure S7:**
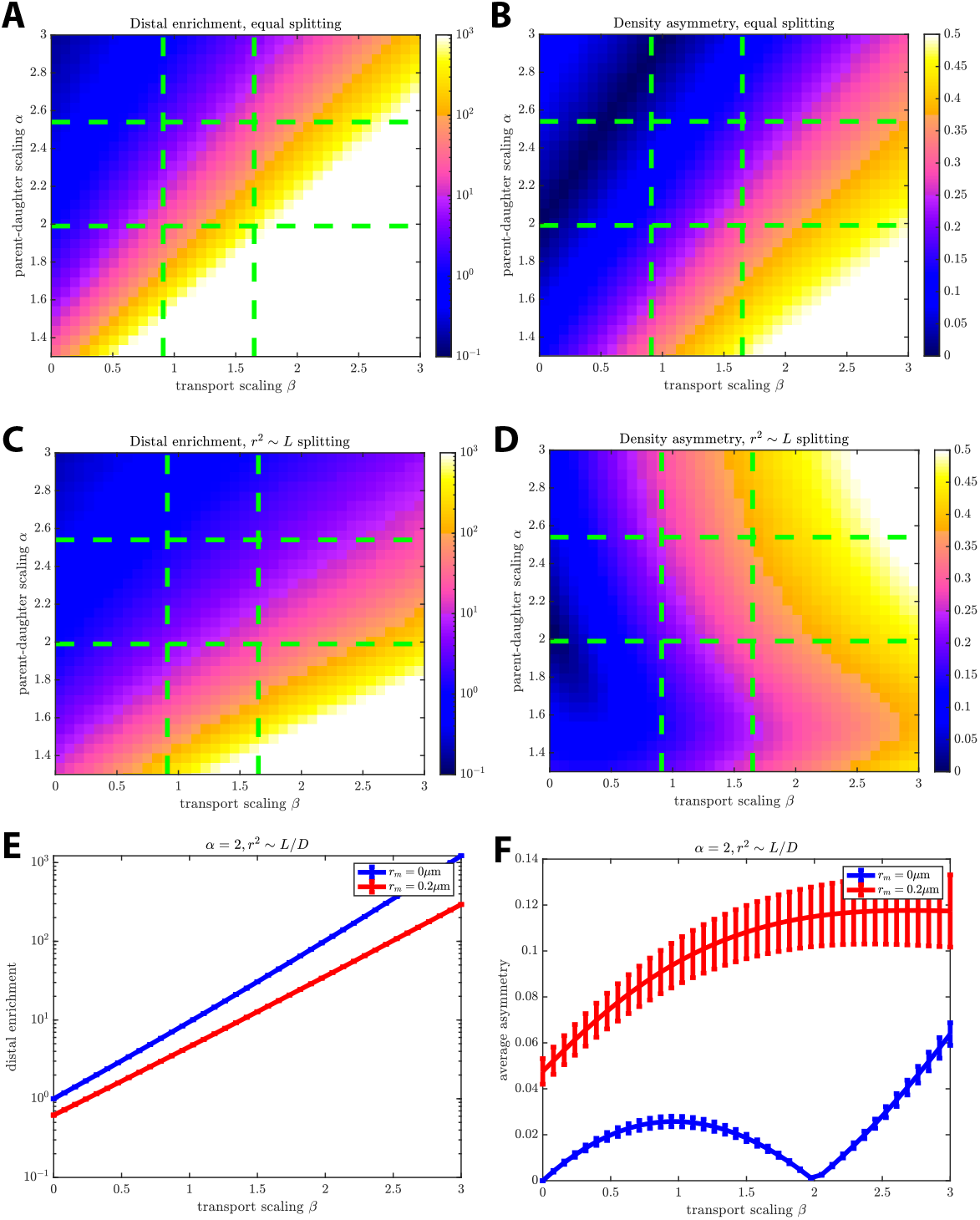
Model mitochondrial distributions in dendrites that obey different scaling rules. A-D: Average mitochondrial distal enrichment (A,C) and density asymmetry (B,D) calculated as a function of α (parent-daughter scaling) and β (transport scaling) for dendrites that obey sister subtree scaling with r_1_=r_2_ (equitable splitting; A-B) or r^2^~L (C-D). Green dashed lines indicate 95% confidence intervals for experimental measurements of α and β. E-F: Average distal enrichment (E) and density asymmetry (F) calculated as a function of β, for r^2^~L/D and parent-daughter scaling with (red) or without (blue) a minimum radius (r_m_): r_0_^2^+r_m_^2^=r_1_^2^+r_2_^2^. Including the minimum radius leads to less distal enrichment (E) and greater density asymmetry (F).

