## Supplemental Methods for "Dendrite architecture determines mitochondrial distribution patterns *in vivo*"

### SUPPLEMENTAL METHODS S1: MEAN-FIELD MODELS FOR MITOCHONDRIAL DISTRIBUTIONS IN A DENDRITIC TREE

**TABLE SM1:** Summary of variables and parameters for mathematical model of mitochondrial density.

**Model parameters**

| Parameter | Description | Expression |
|---|---|---|
| $\alpha$ | parent-daughter branch width scaling rule | $r_0^\alpha = r_1^\alpha + r_2^\alpha$ |
| $\beta$ | scaling exponent of mitochondrial arrest rate vs branch width | $k_s \sim 1/r^\beta$ |
| $k_s$ | mitochondrial arrest rate | |
| $k_w$ | mitochondrial restarting rate | constant |
| $v_i$ | pause-free velocity of mitochondria on branch $i$ | |
| $\mu_0$ | rule for width of first daughter branch (1) relative to parent branch (0) | $\mu_0 = r_1^\alpha / r_0^\alpha$ |
| $r_m$ | minimal radius correction to Da-Vinci parent-daughter scaling | $r_0^2 + r_m^2 = r_1^2 + r_2^2$ |
| $\overline{v}_i$ | average velocity (including) pauses on branch $i$ | |
| $\gamma$ | scaling exponent for average velocity vs branch width | $\overline{v}_i \sim r_i^\gamma$ |

**Model variables**

| Variable | Description | Expression |
|---|---|---|
| $r_i$ | radius of $i^{\text{th}}$ branch | |
| $\ell_i$ | length of $i^{\text{th}}$ branch | |
| $L_0$ | total length of tree with trunk 0 | $L_0 = \sum_{i \in \text{ST}} \ell_i$ |
| $V_0$ | total volume of tree with trunk 0 | $V_0 = \sum_{i \in \text{ST}} \ell_i r_i^2$ |
| $D_0$ | depth of tree with trunk 0 | $D_0 = \ell_0 + \frac{L_1 + L_2}{L_1/D_1 + L_2/D_2}$ |
| $\eta_0$ | relation of volume and trunk area for a tree with trunk 0 | $V_0 = \eta_0 r_0^2$ |
| $\rho_i^{(w)}$ | linear density of motile mitochondria on $i^{\text{th}}$ branch | |
| $\rho_i^{(s)}$ | linear density of stationary mitochondria on $i^{\text{th}}$ branch | |
| $\rho_i$ | total linear density of mitochondria on $i^{\text{th}}$ branch | |
| $c_i^{(w)}$ | volume density of motile mitochondria on $i^{\text{th}}$ branch | $\rho_i^{(w)}/r_i^2$ |
| $c_i^{(s)}$ | volume density of stationary mitochondria on $i^{\text{th}}$ branch | $\rho_i^{(s)}/r_i^2$ |
| $c_i$ | total volume density of mitochondria on $i^{\text{th}}$ branch | |
| $\langle c \rangle_0$ | average mitochondrial volume density over tree with trunk 0 | $\langle c \rangle_0 = \frac{1}{V_0} \sum_{i \in \text{ST}} \ell_i r_i^2 c_i$ |
| $z_0$ | relation between volume density in the trunk and average density in the tree from trunk 0 | $\langle c \rangle_0 = z_0 c_0$ |

#### A. Comparing average subtree densities in models with uniform transport

We first consider models where the mitochondrial transport parameters are spatially uniform (constant velocity $v$ and stopping rate $k_s$ throughout the arbor). Steady state linear densities of motile mitochondria ($\rho_i^w$) and stationary mitochondria ($\rho_i^s$) were computed as described in STAR methods.

#### 1. *Volume densities in arbors obeying Da Vinci parent-daughter scaling*

We consider the case of a tree obeying Da Vinci scaling ($\alpha = 2$) between the parent branch width ($r_0$) and daughter branch widths ($r_1, r_2$), which are related according to $r_0^2 = r_1^2 + r_2^2$. We can solve for the linear densities in the daughter branch as follows:

$$\rho_i = \rho_0 \left( \frac{r_i^2}{r_1^2 + r_2^2} \right) = \rho_0 \frac{r_i^2}{r_0^2}, \quad \text{for } i = 1, 2, \tag{1}$$

where the relationship holds for both the motile and stationary mitochondria densities. The total volume density of mitochondria in a daughter branch $i$ is then

$$c_i = \frac{\rho_i^w + \rho_i^s}{r_i^2} = \frac{\rho_0^w + \rho_0^s}{r_0^2} = c_0 \tag{2}$$

Thus, for the simplest model with uniform mitochondrial transport and conservation of cross-sectional area at each junction (Da Vinci rule), the volume density of mitochondria must be constant throughout the whole tree.

#### 2. *Volume densities in arbors obeying Rall's Law parent-daughter scaling*

Alternately, we can compute the mitochondrial volume density in arbors that obey the Rall's Law relationship between parent and daughter branches (Rule 3, $r_0^\alpha = r_1^\alpha + r_s^\alpha$, with $\alpha = 3/2$). The assumptions of uniform mitochondrial transport (Rule 1 with $\beta = 0$) and mitochondria splitting in proportion to daughter branch area (Rule 2) are still maintained. Here we show that such a model leads to increased mitochonrial densities with distance from the soma, and unequal average densities in asymmetric sister subtrees. The relationships below apply to both motile and stationary mitochondrial densities.

We begin by focusing on a single junction with a parent trunk of radius $r_0$ and linear mitochondrial density $\rho_0$, and daughter trunk radii $r_1, r_2$ and linear densities $\rho_1, \rho_2$. A single parameter $\mu_0 = r_1^\alpha / r_0^\alpha$ describes how the dendritic width is split between sister branches (Rule 4), within the constraints of Rall's law. The fraction of mitochondria that enter the first daughter branch is $\rho_1/\rho_0 = r_1^2/(r_1^2 + r_2^2)$.

The ratio of mitochondrial volume density between the daughter branches and the parent can be written as

$$\frac{c_1}{c_0} = \frac{c_2}{c_0} = \frac{r_0^2}{r_1^2 + r_2^2} = \frac{1}{\mu_0^{2/\alpha} + (1 - \mu_0)^{2/\alpha}} \tag{3}$$

For $\alpha < 2$, this ratio is always above unity ($c_1/c_0 > 1$), except in the edge cases of $\mu_0 = 0$ or $\mu_0 = 1$, which would correspond to one daughter branch disappearing. Thus, the volume density of mitochondria in the daughter branches of a Rall's Law tree is always higher than in the parent branch. This is a direct consequence of the reduced cross-sectional area in the daughter branches.

We next consider whether it is possible to choose values of the sister trunk splitting $\mu_i$ at each junction $i$ in such a way as to ensure equitable mitochondrial distribution in the two sister subtrees. We begin by defining two parameters for a subtree initiating from trunk 0. First, the total volume of the subtree is expressed as $V_0 = \eta_0 r_0^2$. For a symmetric Da Vinci tree, where total cross-sectional area is conserved at each junction, the parameter $\eta_0$ represents the depth of the tree (distance from soma to distal tips). For a symmetric Rall's Law tree, however, the value of $\eta_0$ is less than the depth, due to the narrowing of total cross-sectional area below each junction. Second, we define a parameter $z_0$ relating the average volume density of mitochondria within the subtree, relative to the density within the trunk: $\langle c \rangle_0 = z_0 \rho_0 / r_0^2$. For a Da Vinci tree, under the assumption of uniform mitochondrial transport, $z = 1$ for all junctions regardless of the tree morphology. For a Rall's law tree with at least one junction, the increase in density from parent to daughter branches implies that $z_0 > 1$.

For two sister subtrees, the volume densities in the trunk must be the same ($\rho_1/\rho_2 = r_1^2/r_2^2$, from Rule 1, and therefore $c_1/c_2 = 1$). Consequently, the subtrees will have equal average mitochondrial densities if and only if $z_1 = z_2$. If we want to establish a universal rule for splitting sister trunks (ie: definining $\mu_i$ values at each junction) that depends only on the morphology of the downstream subtree, then the only way to ensure equitable mitochondrial densities throughout a sister subtrees in the arbor would be for all values of $z_i$ to be set to a single constant $z_i = z^*$. For a Rall's Law arbor, we would need to pick a value $z^* > 1$ when setting such a rule. In this case, any subtree consisting of a single long branch (no downstream junctions) would automatically have $z_i = 1$ and would have a lower mitochondrial density than its sister. Furthermore, we show below that if $z^*$ is set close to 1, then the required splitting of branch widths results in most of the tree disappearing and one very long sequence of daughter branches receiving most of the cross-sectional area (Supplemental Figure S4f). Such an extreme rebalancing of branch widths is clearly not representative of tree structure observed *in vivo* and corresponds to an edge case where the arbor is reduced primarily to a single tube rather than a tree. On the other hand, if we set $z^*$ substantially higher than 1, then for many of the junction points it becomes impossible to solve for any value of $\mu$ that would enable equal mitochondrial densities in the sister subtrees.

The two parameters describing the volume and average mitochondrial density in a subtree, can be expressed recursively:

$$\eta_0 = \ell_0 + \eta_1 \mu_0^{2/\alpha} + \eta_2 (1-\mu_0)^{2/\alpha}, \tag{4a}$$

$$z_0 = \frac{\ell_0 + \frac{z_1 \mu_0^{2/\alpha} + z_2 (1-\mu_0)^{2/\alpha}}{\mu_0^{2/\alpha} + (1-\mu_0)^{2/\alpha}}}{\ell_0 + \eta_1 \mu_0^{2/\alpha} + \eta_2 (1-\mu_0)^{2/\alpha}}, \tag{4b}$$

where $\eta_0, z_0$ are the values for a tree with parent trunk 0 and $\eta_{1,2}, z_{1,2}$ are values for the daughter subtrees with trunks 1 and 2.

In Supplemental Figure S4f we consider an example arbor morphology, where the junction connectivities and branch lengths are extracted from a *Drosophila* HS arbor skeleton. Starting from the distal branches of the tree, we recursively solve, where possible, for the value of $\mu_i$ at each junction that would set $z_i = z^*$ for the parent trunk leading to that junction. Where a solution is impossible (always due to the maximum value of $z_i$ being below $z^*$), we pick the splitting that maximizes $z_i$. Red circles in the figure show junctions where a solution was not found that could enable the two sister subtrees to have equal mitochondrial densities. We see that choosing a high value of $z^*$ makes it impossible to enforce equitable mitochondrial densities in many pairs of sister subtrees, in contrast to experimental observations. On the other hand, choosing $z^* \approx 1$ leads to an unrealistic collapse of the arbor to a single primary path in order to maintain equitable mitochondrial distribution.

Overall, these calculations imply that a Rall's tree morphology, together with uniform mitochondrial transport kinetics, leads to increased mitochondrial densities in distal branches but cannot allow for a realistic splitting of branch widths that establishes equal mitochondrial densities between sister subtrees.

#### B. Subtree densities in a Da-Vinci tree with $k_s \sim 1/r^2$

Rather than assuming spatially constant mitochondrial motility, an alternative model can be constructed where mitochondria are more likely to halt on narrower branches, while the restarting rate $k_w$ and pause-free velocities $v$ remain constant. One simple model for width-dependent stopping would be to set the rate inversely proportional to the cross-sectional area of each branch: $k_{s,i} = k_s^*/r_i^2$, corresponding to $\beta = 2$ in scaling Rule 3. We then consider the distribution of stopped mitochondria in different dendritic branches. With $k_s \sim 1/r^2$, the linear density of stationary mitochondria in branch $i$ is given by

$$\rho_i^{(s)} = \frac{k_s^*}{k_w} \rho_i^{(w)} / r_i^2 \tag{5}$$

Where $\rho_i^{(w)} = \rho_{+,i} + \rho_{-,i}$ is the motile linear density of mitochondria. At a junction with daughter branches $1, 2$, this motile linear density splits according to $\frac{\rho_1^{(w)}}{\rho_2^{(w)}} = \frac{r_1^2}{r_2^2}$.

##### 1. *Comparing average densities in sister subtrees*

In a tree with Da Vinci scaling, the volume density of motile mitochondria is spatially constant, so that all branches have $c_i^{(w)} = c_{\text{trunk}}^{(w)} \rho_{\text{trunk}}^{(w)} / r_{\text{trunk}}^2$. We can then calculate the average volume density of the stationary population in a subtree with total volume $V_{\text{ST}}$ and total branch length $L_{\text{ST}}$:

$$\left\langle c^{(s)} \right\rangle_{\text{ST}} = \frac{\sum_{i \in ST} \rho_i^{(s)} \ell_i}{\sum_{i \in ST} r_i^2 \ell_i} = \frac{\frac{k_s^*}{k_w} c_{\text{trunk}}^{(w)} \sum_i \ell_i}{V_{ST}} \sim \frac{L_{ST}}{V_{ST}}, \tag{6}$$

where the summations are over all branches in the subtree.

Therefore, the ratio between the average stopped mitochondrial densities in sister subtrees becomes:

$$\frac{\left\langle c^{(s)} \right\rangle_1}{\left\langle c^{(s)} \right\rangle_2} = \frac{L_1/V_1}{L_2/V_2} \tag{7}$$

Keeping in mind that $\langle c \rangle = \left\langle c^{(w)} \right\rangle + \left\langle c^{(s)} \right\rangle$ and that $c^{(w)}$ is the same for all branches in a Da Vinci tree, we see that equitable distribution of mitochondria between sister subtrees can be achieved only if the volume of each sister subtree is proportional to its total length:

$$\frac{L_1}{V_1} = \frac{L_2}{V_2}. \tag{8}$$

Notably, increased stopping of mitochondria in narrower branches implies that more distal sections of the dendritic tree will tend to have higher mitochondrial densities. We therefore predict that the combination of Da Vinci scaling of branch widths ($r_0^2 = r_1^2 + r_2^2$) together the the proportionality of subtree length and total volume (Eq. 8) will give rise to mitochondrial densities that both increase with distance from the soma and are equal between sister subtrees, as observed for experimental data.

*2. Sister branch radii splitting for $L \sim V$ relationship*

For a Da Vinci arbor, the proportional relationship between sister subtree length and volume (Eq. 8) can be achieved via a particular morphological rule governing the relative trunk widths of sister substrees emerging from the same junction (Rule 4).

We begin by defining the depth of a tree ($D$) via a recursive approach. For a subtree consisting of a single branch of length $\ell_1$, the depth is simply defined as that branch length ($D_1 = \ell_1$). Next, we consider a tree with trunk of index 0, splitting at a downstream junction between subtree trunks 1 and 2. We define the depth of the tree according to the following formula:

$$D_0 = \ell_0 + \frac{L_1 + L_2}{L_1/D_1 + L_2/D_2}, \tag{9}$$

where $D_1, D_2$ are the depths and $L_1, L_2$ are the total branch lengths of the subtrees starting with branch 1 and 2, respectively. Conceptually, this expression averages the inverse depths of the two subtrees, weighted by their respective lengths, and adds on the length of the parent trunk. We note that in the case where the two subtrees have the same depth ($D_1 = D_2$) then the overall depth of the tree becomes $D_0 = \ell_0 + D_1$. Thus, in an arbor where all distal tips are the same distance from the parent node, the depth of the tree will simply be equal to that distance.

We now consider the specific case of a Da Vinci arbor that additionally obeys the criterion in Eq. 8, where the volume of a sister subtree is proportional to its total length. As before, we express the volume of the arbor in terms of the prefactor $\eta_0$ according to $V = \eta_0 r_0^2$. We then show by induction that under these assumptions the prefactor is equal to the depth: $D_0 = \eta_0$. First, we use the length-volume proportionality to express the volume of each subtree in terms of the parent volume and the subtree lengths according to:

$$\begin{aligned} &V_0 = \ell_0 r_0^2 + V_1 + V_2, \\ &V_i = \frac{L_i}{L_1 + L_2}(V_1 + V_2) = \frac{L_i}{L_1 + L_2}\left(V_0 - \ell_0 r_0^2\right) = \eta_i r_i^2, \quad i = 1, 2 \end{aligned} \tag{10}$$

Next, we can apply the Da Vinci law relating parent and daugher branch widths:

$$\begin{aligned} r_0^2 = r_1^2 + r_2^2 &= \frac{L_1/\eta_1 + L_2/\eta_2}{L_1 + L_2}\left(V_0 - \ell_0 r_0^2\right), \\ V_0 &= \left(\ell_0 + \frac{L_1 + L_2}{L_1/\eta_1 + L_2/\eta_2}\right) r_0^2 = \eta_0 r_0^2 \end{aligned} \tag{11}$$

Thus, we see that if the two subtrees have depths $D_1 = \eta_1$ and $D_2 = \eta_2$, then the overall tree will also have $D_0 = \eta_0$, where depth is defined according to Eq. 9. Since single branches have $\eta_i = D_i$ by definition, this argument implies that all trees obeying Da Vinci scaling and length-volume proportionality, have volume given by $V_0 = D_0 r_0^2$.

Finally, we note that, for a Da Vinci tree, the proportionally of length and volume can now be translated directly into a relationship between sister subtree trunk widths:

$$\begin{aligned} \frac{L_1/V_1}{L_2/V_2} &= \frac{L_1/(D_1 r_1^2)}{L_2/(D_2 r_2^2)} = 1, \\ \frac{r_1^2}{r_2^2} &= \frac{L_1/D_1}{L_2/D_2} = \frac{b_1}{b_2} \end{aligned} \tag{12}$$

where we define the 'bushiness' of a subtree ($b_i$) as its total length divided by its depth: $b_i = L_i/D_i$. Trees with high bushiness are broader, in the sense of having a greater total length of branches at a given depth, arising from more frequent junctions (Figure 4b).

Overall, we have shown that in an arbor obeying the Da Vinci rule ($\alpha = 2$), where mitochondrial stopping is inversely proportional to branch area ($\beta = 2$), equal densities of mitochondria between sister subtrees will be obtained if the sister trunk areas are split in proportion to the subtree bushiness (Eq. 12).

#### C. Average subtree densities for general transport behavior in Da Vinci arbors

We next consider a generalization of the mitochondrial distribution model to the case where both the stopping rate $k_{s,i}$ and the pause-free velocity $v_i$ can vary depending on the branch width. At steady state, the conservation of incoming and outgoing flux into a branch junction gives a relationship between the motile mitochondria density $\rho_0^{(w)}$ in the parent trunk 0 and the daughter branches $1, 2$. We maintain the assumption that the splitting of mitochondrial flux into each daughter branch is proportional to the cross-sectional area. Specifically, this gives the two conditions:

$$v_o \rho_o^{(w)} = v_1 \rho_1^{(w)} + v_2 \rho_2^{(w)} \tag{13a}$$

$$\frac{v_1 \rho_1^{(w)}}{v_2 \rho_2^{(w)}} = \frac{r_1^2}{r_2^2}. \tag{13b}$$

The density of stationary mitochondria in each branch is given by $\rho_i^{(s)} = \frac{k_{s,i}}{k_w} \rho_i^{(w)}$.

Assuming a Da Vinci relationship between parent and daughter branch widths, we can solve for the volume density of mitochondria in daughter branches as follows:

$$\begin{aligned} v_0 \rho_0^{(w)} &= v_1 \rho_1^{(w)} \left(1 + \frac{r_2^2}{r_1^2}\right) = \frac{v_1 \rho_1^{(w)} r_0^2}{r_1^2} \\ c_1^{(w)} &= \rho_1 / r_1^2 = c_0^{(w)} \frac{v_0}{v_1}. \end{aligned} \tag{14}$$

Consequently, throughout the entire arbor, the motile volume density in each branch can be written in terms of the local velocity and the density in the parent trunk of the full tree: $c_i^{(w)} = c_{\text{trunk}}^{(w)} \frac{v_{\text{trunk}}}{v_i}$. The total volume density on a branch, including motile and stationary mitochondria, can be expressed in terms of the average velocity (with pauses included), given by $\overline{v}_i = \frac{k_w}{k_w + k_{s,i}} v_i$. Specifically:

$$c_i = c_i^{(w)} + c_i^{(s)} = \left(\frac{k_{s,i} + k_w}{k_w}\right) c_{\text{trunk}}^{(w)} \frac{v_{\text{trunk}}}{v_i} = c_{\text{trunk}}^{(w)} v_{\text{trunk}} / \overline{v}_i \tag{15}$$

The average volume density of mitochondria in a subtree is then given by

$$\langle c \rangle_{\text{ST}} = \frac{\sum_{i \in \text{ST}} c_i^{(w)} r_i^2 \ell_i}{\sum_{i \in \text{ST}} r_i^2 \ell_i} = \frac{1}{V_{\text{ST}}} \sum_{i \in \text{ST}} \frac{r_i^2 \ell_i c_{\text{trunk}}^{(w)} v_{\text{trunk}}}{\overline{v}_i} = c_{\text{trunk}}^{(w)} v_{\text{trunk}} \left\langle \frac{1}{\overline{v}} \right\rangle_V, \tag{16}$$

where the final term denotes the volume-weighted average of the inverse velocity over the subtree: $\left\langle \frac{1}{\overline{v}} \right\rangle_V = \left( \sum_{i \in \text{ST}} r_i^2 \ell_i (1/\overline{v}_i) \right) / V_{\text{ST}}$.

We consider the case where the average velocity (including pauses) along a branch scales as a power law of the branch width: $\overline{v}_i \sim r_i^\gamma$. Under this assumption, the ratio of sister subtree densities is given by

$$\frac{\langle c\rangle_1}{\langle c\rangle_2} = \frac{\left\langle \frac{1}{\bar{v}} \right\rangle_{V_1}}{\left\langle \frac{1}{\bar{v}} \right\rangle_{V_2}} = \frac{V_2}{V_1} \frac{\sum_{i\in \mathrm{ST}_1} r_i^{2-\gamma}\ell_i}{\sum_{i\in \mathrm{ST}_2} r_i^{2-\gamma}\ell_i}. \tag{17}$$

In the case that $\gamma = 2$, this relationship reduces to $\langle c\rangle_1 / \langle c\rangle_2 = (L_1/V_1)/(L_2/V_2)$, and equal densities of mitochondria between sister subtrees are again achieved when the subtree volume is proportional to its total length. A particular case that leads to $\gamma = 2$ is where restarting rates are low ($k_{s,i} \gg k_w$ throughout most of the tree), pause-free velocities are constant, and the stopping rate scales inversely with cross-sectional area ($k_{s,i} \sim 1/r_i^2$). This is the simplified case considered in the main text.
